# Mechanism Study on the Effect of Kidney Tonifying and Liver Soothing Methods on Mouse Oocyte Quality by Regulating OSFs and Smads Pathways

**DOI:** 10.1101/2024.02.12.579988

**Authors:** Ruijuan Zhang, Rui Jia, Jing Bai, Huilan Du, Liyun Yang

**Affiliations:** Hebei University of Chinese Medicine, 050200;2. Langfang Vocational College of Health, 065001

**Keywords:** kidney tonifying method, Liver soothing method, oocyte, BMP-6, ALK-2, ALK-6, Smad1, Smad5, Smad8, Smad4

## Abstract

Objective: To observe the effects of kidney tonifying and liver soothing methods on the secretion of factor BMP-6, related receptors ALK-2/6, and downstream Smads pathway Smad1/5/844 in oocytes, and to explore the targeted mechanisms of their effects on mouse oocyte quality; Method: Healthy female mice aged 6-7 weeks were randomly divided into 6 groups, namely high and low dose kidney tonifying groups; High – and low-dose liver soothing groups; Control group and normal group. The high-dose and low-dose groups of tonifying the kidney were given oral solution of tonifying the kidney and regulating the meridian at 5.4g/ml and 2.7g/ml, respectively. The high-dose and low-dose groups of soothing the liver were given suspension of Xiaoyao Pill at 0.6g/ml and 0.3g/ml, respectively. The control group and normal group were given distilled water by gavage. Immunohistochemistry and Western blot were used to detect the expression of BMP-6, ALK-2/6, and Smad1/5/8/4 proteins in oocytes, while PCR was used to detect the mRNA expression of these indicators in oocytes; Result: Both methods can increase the expression of BMP-6 in oocytes of mice in the treatment group, activate ALK-2/6, and phosphorylate Smad1/5/8 to bind with Smad4, initiating signal transduction; The high-dose group of kidney tonifying is superior to other groups in regulating BMP-6, ALK-2/6, and Smad1/5/4. Conclusion: Kidney tonifying and liver soothing methods can regulate BMP-6 and its Smads pathway through different mechanisms to improve mouse oocyte quality.

## 1. Introduction

Due to changes in lifestyle and environmental pollution, infertility has become a key factor that seriously affects human reproductive health and family and social harmony of couples of childbearing age. Its incidence rate is increasing, which is about 8% –17% in China, of which ovarian factors account for 25% –35% [1]. With the rapid development of assisted reproductive technology (ART) in humans, in vitro fertilization embryo transfer (IVF-ET)[2] has a success rate of over 90% in follicle recruitment and oocyte acquisition. The quality of embryos has been significantly improved, and the number of high-quality embryos obtained has also increased significantly. However, its clinical pregnancy rate has not been satisfactory. In recent years, traditional Chinese medicine has made significant progress in improving oocyte quality, inducing ovulation, increasing pregnancy rates, and reducing the toxic side effects of Western medicine [3,4,5,6]. Our research group has previously studied the effects of kidney tonifying and liver soothing methods on the reproductive function of anovulatory model rats. It has been confirmed that kidney tonifying and liver soothing methods can improve the morphology of the pituitary gland and ovaries, regulate the levels of neurotransmitters in the hypothalamus and reproductive endocrine hormones, promote follicular development and maturation, and promote uterine development. They can also regulate the endometrial environment to facilitate the implantation of fertilized eggs[7]; We studied the effects of Bushen Tiao Jing Fang and Xiaoyao San on the pregnancy outcomes and OSFs of IVF-ET, and confirmed that the addition of Bushen Tiao Jing Fang and Xiaoyao San can increase the number of eggs obtained, improve the rate of high-quality embryos and clinical pregnancy rates [8]. However, the mechanism by which they affect ovulation is not yet clear. On the basis of previous research, this project aims to further investigate the mechanism of the effects of kidney tonifying and liver soothing methods on oocyte development through animal experiments, laying a theoretical foundation for the clinical combination of IVF-ET treatment for infertility.

## 2. Materials and Methods

### 2.1 Materials

#### 2.1.1 Experimental animals

600 healthy female Kunming mice aged 6-7 weeks, weighing 28-32g, clean grade, purchased from Hebei Experimental Animal Center, certificate number: 1208011. Free drinking and eating, keeping the temperature in the breeding room at (20 ± 1) ℃, relative humidity of about 50%, automatic control of light for 12 hours, and darkness for 12 hours.

#### 2.1.2 Experimental drugs

Injection of PMSG (batch numbers 20120410, 20120716), Shanghai Institute of Family Planning Science; HCG for injection (batch numbers 120108, 120502, 120705), produced by Lizhu Pharmaceutical Factory of Lizhu Group; Xiaoyao Pills (batch number 111133), purchased from Wanxi Pharmaceutical Co., Ltd. in Henan Province, from Lerentang Pharmacy in Shijiazhuang City (200 and 100 Xiaoyao Pills were respectively dissolved in 125ml distilled water to form suspensions, with high-dose raw materials of 0.6g/ml and low-dose raw materials of 0.3g/ml); The prescription for tonifying the kidney and regulating the meridians (consisting of Rehmannia glutinosa, Cornus officinalis, Chinese yam, Lycium barbarum, Ligustrum lucidum, Ziheche, Epimedium, Cuscuta chinensis, Raspberry, Rhizoma Cyperi, Radix Paeoniae Alba, Angelica Sinensis, etc.), identified as authentic by the Department of Traditional Chinese Medicine of Hebei Medical University, is made into oral liquid by the Department of Traditional Chinese Medicine of the university (high dosage crude drug 5.4g/ml, low dosage crude drug 2.7g/ml); Hyaluronidase: H8030, Solarbio, Beijing; 0.01mol/L PBS working solution (pH 7.2-7.4): NaCl8.0g, KCl0.2g, stored at 4 ℃; Primer synthesis: Bioengineering (Shanghai) Co., Ltd; RNA extraction kit: ABI, USA; M-MLV reverse transcription kit: Promega, USA; SYBR Green I Fluorescence Quantitative PCR Kit: Females, Canada;

Immunohistochemistry:Firstantibody:BMP-6:bs-3671R;ALK-6:sc-25455;Smad1: bs-1619R;Smad5:bs-2973R;Smad8:bs-4253R;Smad4:bs-0585R;DABdyeingSolution, BeijingBoosenBiotechnology Co., LTD.; ALK-2: PAB3474, Abnova Corporation; Second antibody: sp-9001, Beijing Zhongshan Jinqiao Biotechnology Co., LTD.

Western blot:First antibody: BMP-6: sc-7406; ALK-6: sc-5679,Santa Cruz Corporation; ALK-2: PAB3474, Abnova Corporation; Smad1:1649-1; Smad4:1676-1; Smad5:1682-1, Epitomics Corporation; Smad8: bs-4253R, Beijing Boosen Biotechnology Co., LTD. Rabbit secondary antibody: 611-131-122; Sheep second antibody: 605-731-125, Rockland;ß-actin: R1207-1, Hangzhou Hua’an Biotechnology Co., LTD.; Coomassie Bright Blue Dye Solution for protein quantification: PA102, Beijing TIANGEN Company;

#### 2.1.3 Main instruments

Continuous zoom microscope: XTL-1 type, Beijing Electronic Optics Equipment Factory; Continuous zoom microscope: SMZ-T4 type, Chongqing Aote Optical Instrument Co., Ltd; OLYMPUS upright microscope: BX51T-PHD-J11; Camera: Nikon D60, Japan; Electric constant temperature water bath: H W21. Cu600 type, Shanghai Yiheng Technology Co., Ltd; Electronic balance: SARTORIUSAG, Germany; Egg stripping needle: Shenzhen Huanhao Technology Co., Ltd; Glass slide: Qingdao, China; Micropipette: Eppendorf, USA; Vortex mixer: VORTEX-5 type, Haimen Qilinbei Instrument Manufacturing Co., Ltd; Ice maker: AF10 SCOTSMAN from the United States; UV spectrophotometer: ND2000 model, Thermo, USA; Fluorescence quantitative PCR System: ABI 7300, American company ABI; Low temperature centrifuge: RS-28 type, Hercules AG, Germany; Versa Max TM ELISA reader: Molecular Device, USA; Electrophoresis tank: DYCZ-21 type, Beijing Liuyi Instrument Factory; Electrophoresis apparatus; Semi dry protein imprinting transfer tank: Bio Rad, USA; PVPF membrane: Millipore, USA; Odyssey Infrared Fluorescence Scanning Imaging System: LI-COR, USA

### 2.2 Experimental methods

#### 2.2.1 Grouping

After adaptive feeding of mice for one week, they were completely randomized into 6 groups using a random number table method, including high-dose kidney tonifying group, low-dose kidney tonifying group, high-dose liver soothing group, low-dose liver soothing group, control group, and normal group, with 100 mice in each group.

#### 2.2.2 Method of administration

Each group of mice was divided into the following groups: the high-dose and low-dose groups of kidney tonifying were given oral solution of kidney tonifying and meridian regulating formula, the high-dose and low-dose groups of liver soothing were given suspension of Xiaoyao Pill, and the COH group and blank control group were given distilled water by gavage. At 8:00 every day, each group of mice was orally administered with 1ml/100g body weight for 10 consecutive days. On the 11th day, the high-dose group of kidney tonifying, low-dose group of kidney tonifying, high-dose group of liver soothing, low-dose group of liver soothing, and COH group were simultaneously intraperitoneally injected with PMSG 10 IU per animal. After 48 hours, HCG 10 IU per animal was intraperitoneally injected [9]; The blank control group received daily vaginal smears and observed the estrous cycle.

#### 2.2.3 Sample collection

The normal group of mice showed a large amount of keratinized epithelial cells on vaginal smears the next day, while the other groups of mice were euthanized by cervical spondylosis 16 hours after injection of HCG. The abdominal cavity of the experimental mice was surgically opened in an ultra clean workbench, and both fallopian tubes were taken out and placed in sterile culture dishes. Under a stereomicroscope, a 1ml syringe needle was used to cut through the transparent and enlarged ampulla of the fallopian tubes, and flocculent substances were observed to be released, which are called cumulus oocyte complexes (COCs), Digest granulosa cells with hyaluronidase, wash repeatedly with PBS to obtain M Ⅱ stage oocytes, partially store at –80 ℃ for Western blot, and partially use for immunohistochemistry.

#### 2.2.4 Detection method

##### 2.2.4.1 Immunohistochemical detection of the expression of BMP-6, ALK-2/6, Smad1/5/8/4 proteins in mouse oocytes in each group

Xylene is dewaxed into water, followed by gradient alcohol dehydration. Incubate at room temperature with 3% hydrogen peroxide for 10 minutes, rinse with PBS for 5 minutes × 3 times, seal with sealing solution for 30-40 minutes, remove excess sealing solution, add primary antibody (working concentration 1:50) dropwise, incubate at 37 ℃ in a wet box for 60 minutes, and rinse with PBS for 5 minutes × 3 times; Add biotin labeled secondary antibody dropwise, incubate at room temperature for 10 minutes, and rinse with PBS for 5 minutes × 3 times; Add streptavidin peroxidase solution dropwise, incubate at room temperature for 10 minutes, and rinse with PBS for 5 minutes × 3 times, DAB staining, hematoxylin re staining of cell nucleus, differentiation with 1% hydrochloric acid ethanol, ammonia water returning to blue, rinsing with double distilled water, air drying, neutral gum sealing. The IHS score [10] was used to observe the results by combining the percentage of positive cells and the strength of positive cell staining under the light microscope.

##### 2.2.4.2 Real-Time RT-PCR was used to detect the expression of BMP-6, ALK-2/6, Smad1/5/8/4 mRNA in mouse oocytes in each group

(1) Primer design:(table1)
(2) RNA extraction: Put the oocytes in the cryogenic centrifuge for agarose gel electrophoresis detection, and operate according to the instructions.
(3) RNA purity measurement: OD260 and OD280 were measured using an ND2000 UV spectrophotometer, and the OD260/OD280 ratio was calculated to be between 1.8-2.0. This indicates that the extracted RNA has high purity, minimal contamination, and can be used for subsequent reverse transcription reactions.
(4) Reverse Transcription (RT)(table 2):
(5) Fluorescence quantitative PCR detection: The composition of the PCR reaction solution (table 3)
(6) Real-time fluorescence quantitative PCR result analysis: After amplification, enter the SDS V1.3 software result analysis interface for analysis.

##### 2.2.4.3 Western blot was used to detect the expression of BMP-6, ALK-2/6, Smad1/5/8/4 proteins in mouse oocytes from each group

Collect oocytes from each group, resuspend them in cell lysate, and use Bradford method for protein quantification (follow the instructions). SDS polyacrylamide gel electrophoresis was used to analyze the protein composition of the samples. After that, the gel was semi dried, and then the PVDF membrane was taken out and placed in TBST sealing solution containing 5% skimmed milk powder, and sealed at 37 ℃ for 2h. Then place it in appropriately diluted primary and secondary antibody solutions. Odyssey luminescence instrument detects antibody specific binding bands.

**Table 1.**
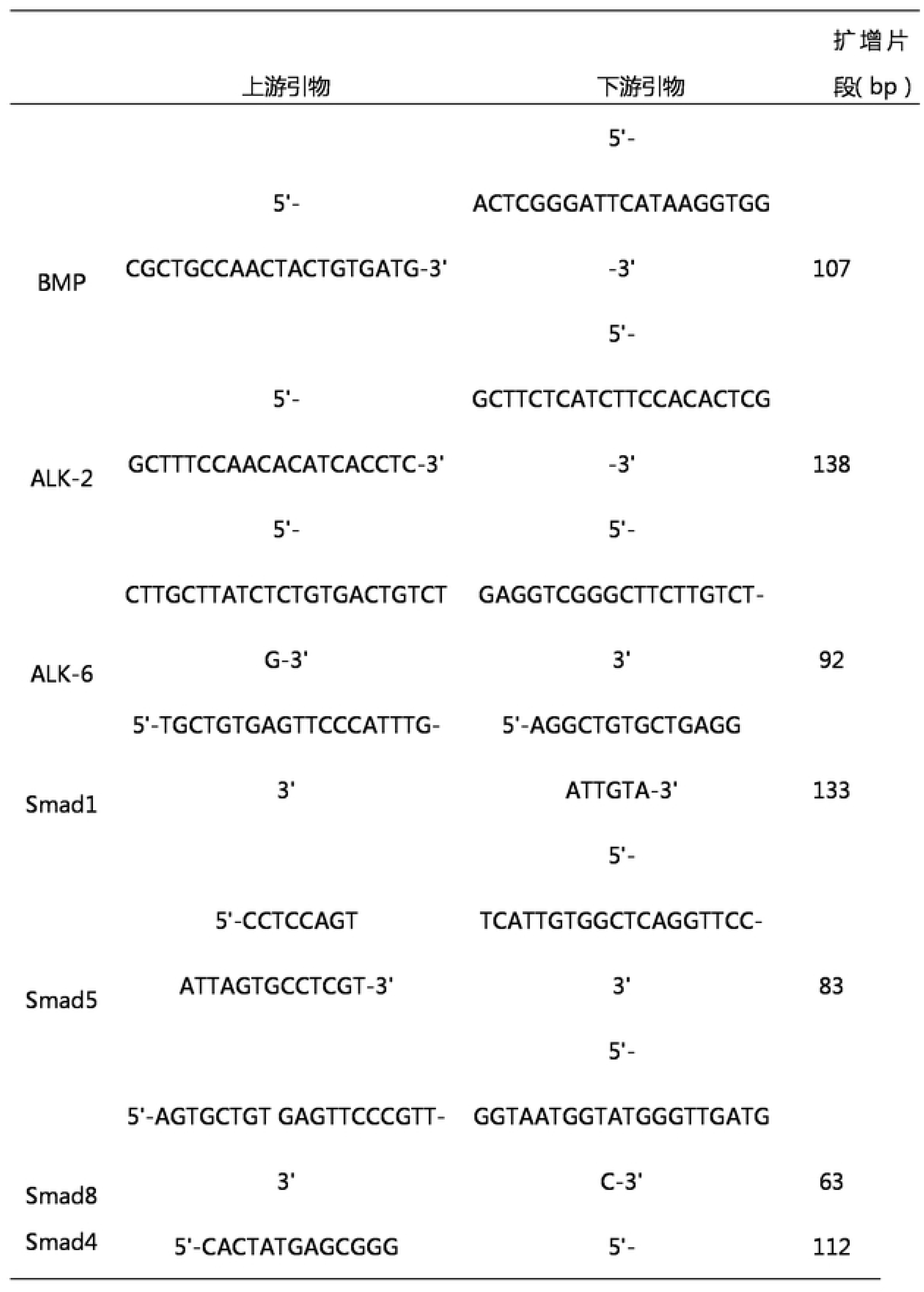
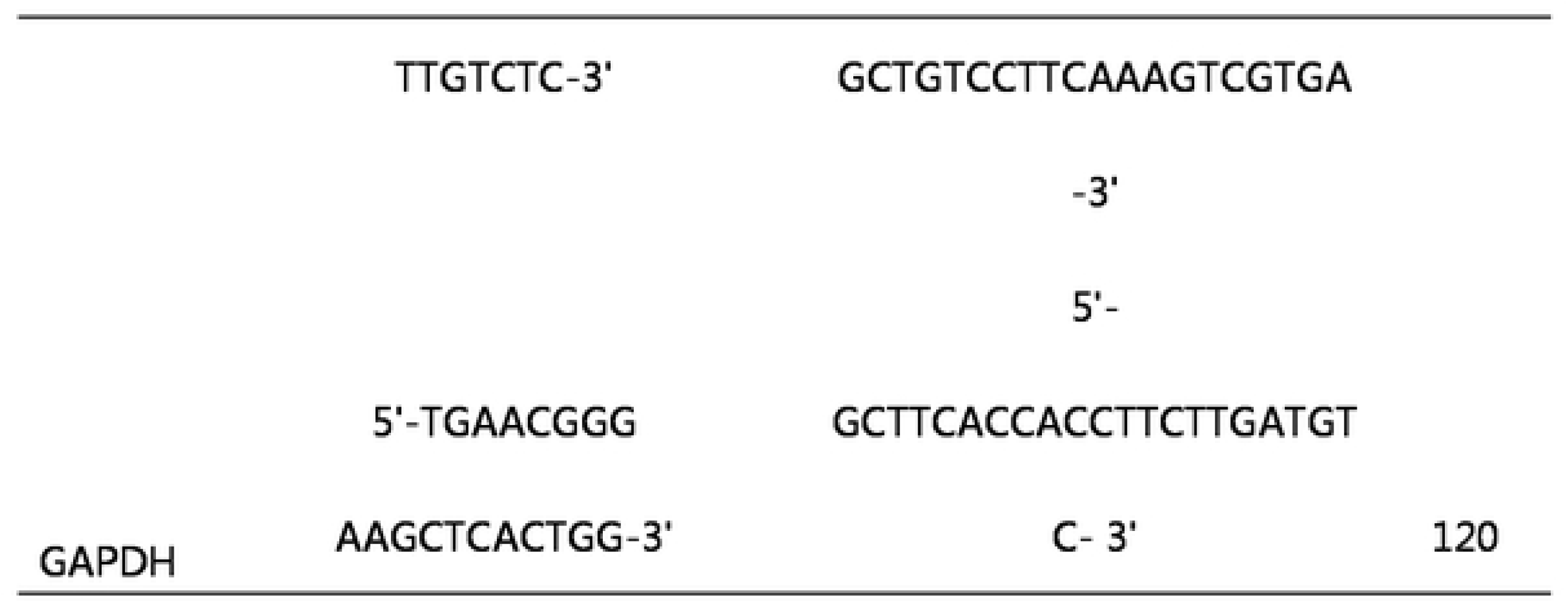
primer design.

**Table 2.** The composition of the reverse transcription reaction liquid.

| Component | Volume (μl) |
| --- | --- |
| 5 × buffer | 4 |
| dNTP | 2 |
| Rnasin | 1 |
| RNA | 5 |
| Random Primer | 1 |
| M-MLV | 1 |
| Nucleasr-Free Water | 6 |
| Total | 20 |

Reverse transcription at 42 °C for 50 min and inactivation of reverse transcriptase at 95 °C for 5 min.

**Table 3.** The composition of the PCR reaction solution.

| Component | Volume ( μl ) |
| --- | --- |
| R.T.product ( cDNA ) | 1 |
| 2×Master Mix | 25 |
| Upstream primers | 1 |
| Downstream primers | 1 |
| Deionized water | 22 |
| Total | 50 |

PCR thermal cycle parameters: 95 °C for 5min, followed by a three step reaction: 94 °C for 30s, 58 °C for 30s, and 72 °C for 30s. Forty cycles were conducted, and fluorescence signals were collected at the third step of each cycle, namely, 72 °C for 30s.

### 2.3 Statistical methods

All experimental data were statistically processed using the SPSS 21.0 software package. Quantitative data is expressed as mean ± standard deviation (x ± S). If normality and homogeneity of variance are met, one-way ANOVA is used for comparison between multiple groups. Otherwise, rank sum test is used; The count data was analyzed using X2 test, and P<0.05 indicates a statistically significant difference.

## 3. Results

### 3.1 General situation of animals (Figures 1 to 4)

The mice in each group developed well, with white and glossy fur, good mental state, sensitive reactions, normal diet and bowel movements, and increased weight. After removing the fallopian tubes from the surgical open abdominal mice, an enlarged and transparent ampulla of the fallopian tubes can be seen, which is filled with flocculent cumulus oocyte complexes. Piercing the ampulla of the fallopian tubes can release the cumulus oocyte complex, which is radiating

**Fig. 1.**
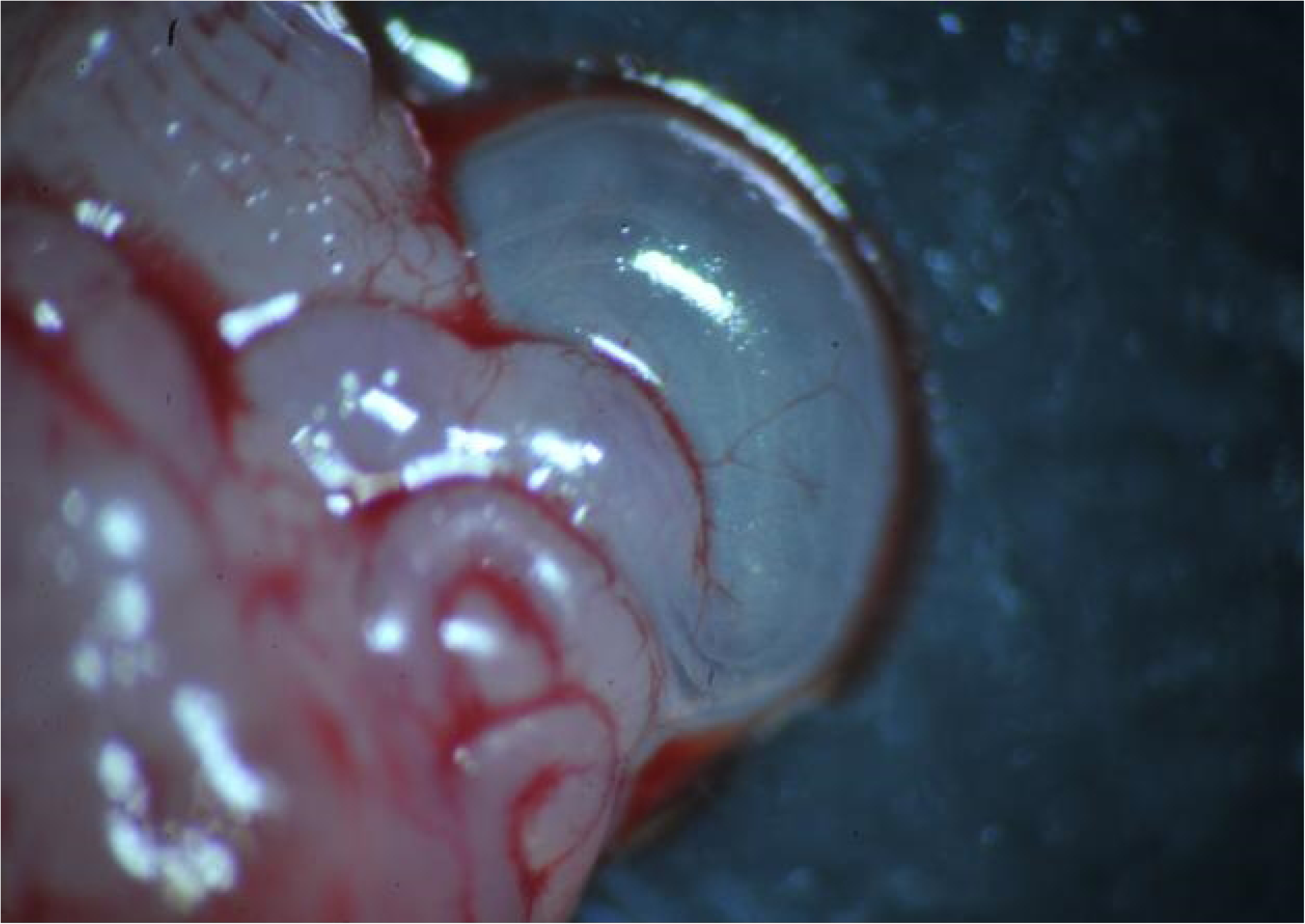
Inflated ampulla of oviduct(×40)

**Fig. 2.**
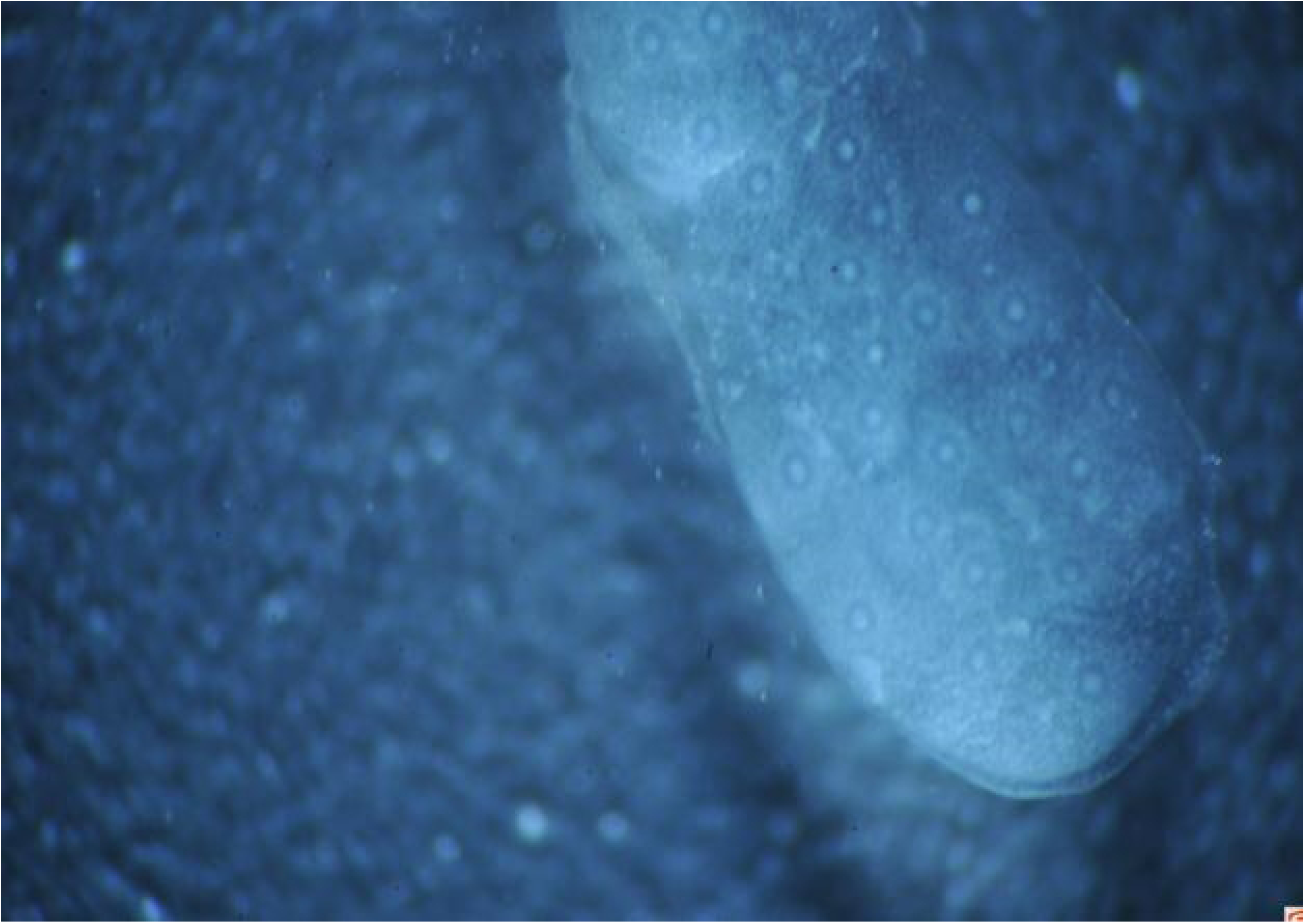
The cumulus oocyte complexes released from oviduct(×40)

**Fig. 3.**
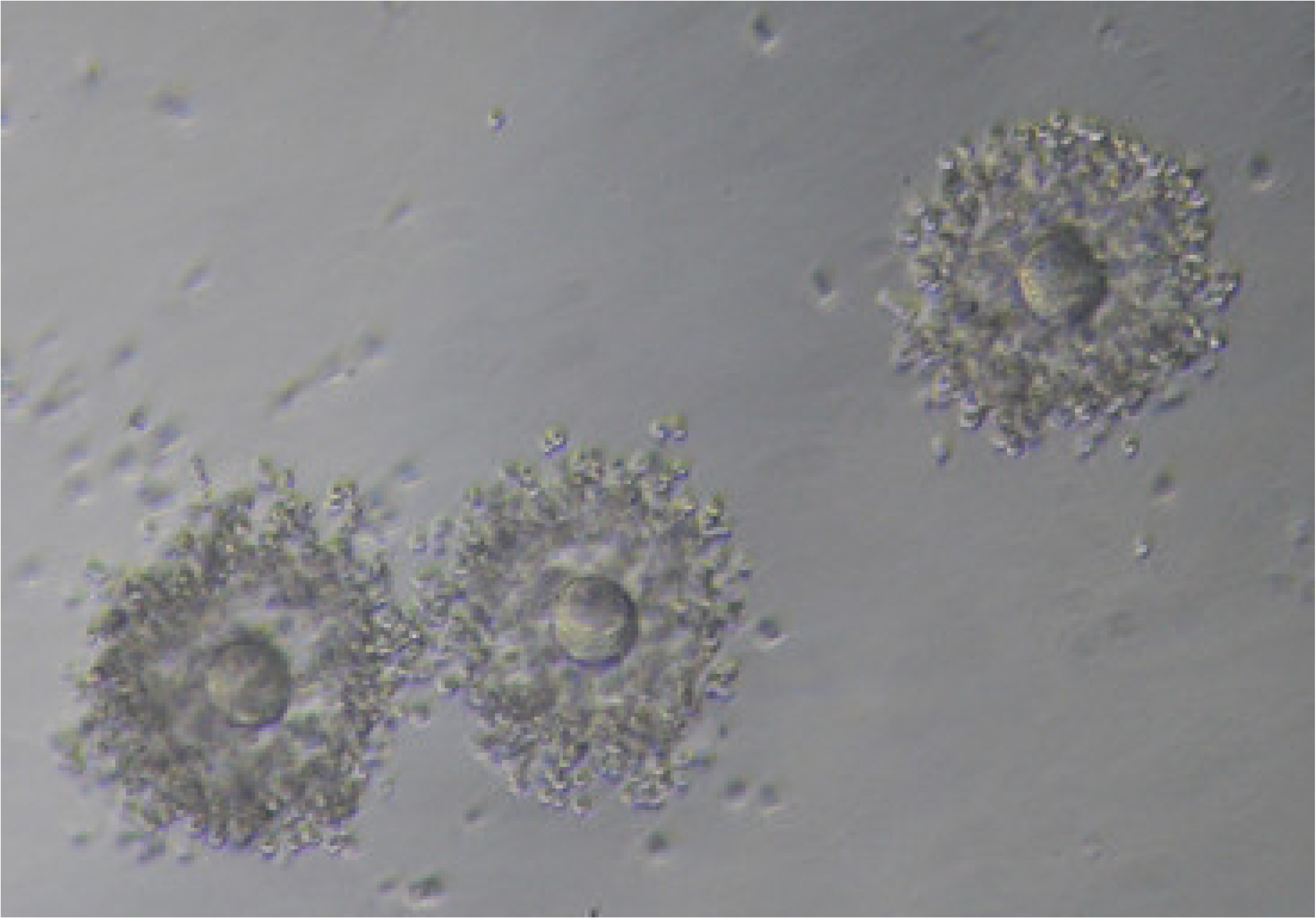
MⅡoocytes(×100)

**Fig. 4.**
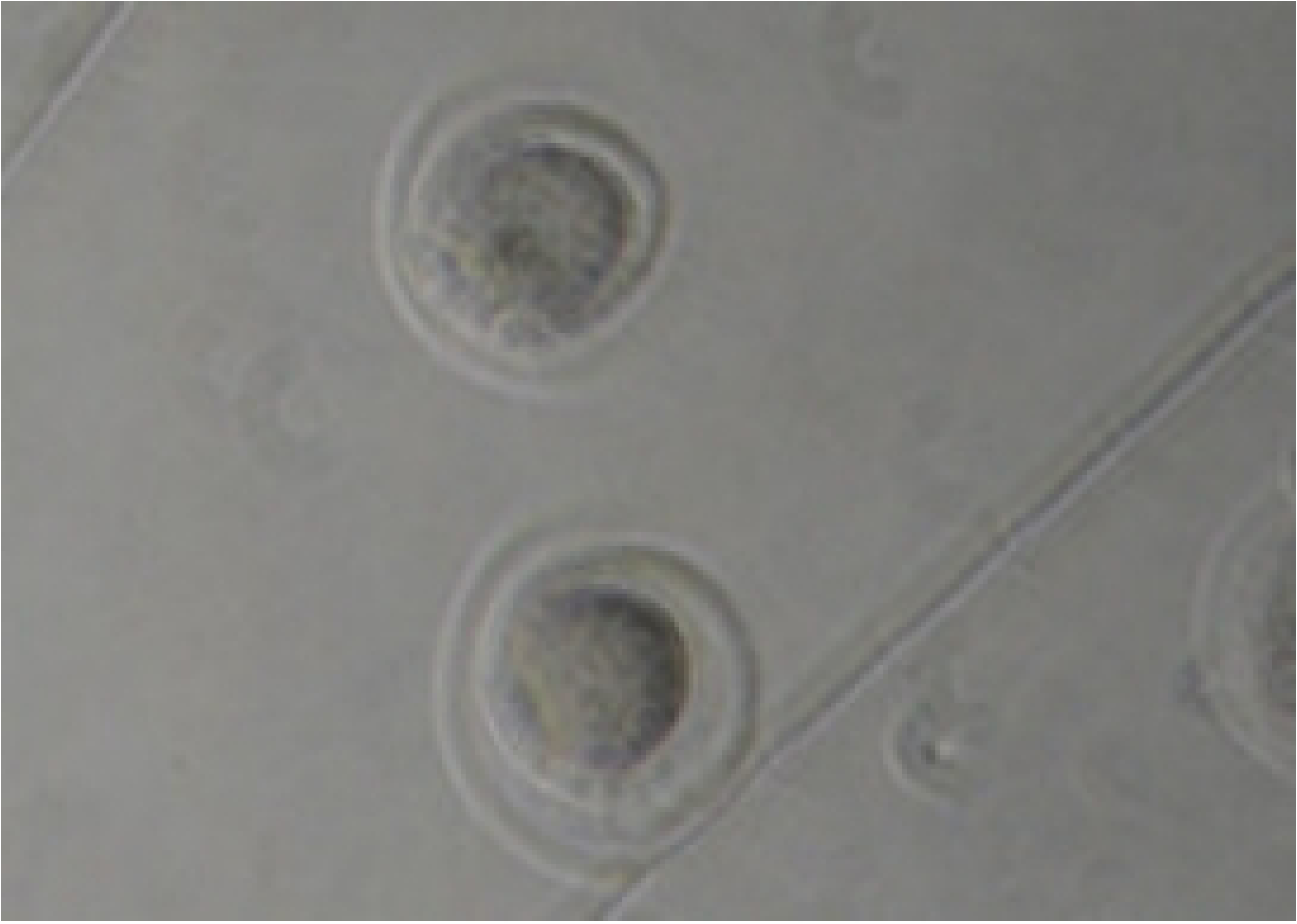
MⅡ oocytes(×400)

### 3.2 Comparison of immunohistochemical methods for detecting protein expression of BMP-6, ALK-2/6, Smad1/5/8/4 in mouse oocytes from different groups

#### 3.2.1 Comparison of BMP-6 protein expression (Figure 5-7, Table 4)

There was no statistically significant difference (P>0.05) between the blank control group and the COH group; The IHS scores of each treatment group were higher than those of the blank control group and COH group (P<0.05); The expression in the high-dose group of kidney tonifying was better than that in the low-dose group of kidney tonifying, high-dose group of liver soothing, and low-dose group (P<0.05); The IHS score of the high-dose liver soothing group increased compared to the low-dose group (P<0.05); There was no statistically significant difference between the low-dose groups (P>0.05)

**Fig. 5.**
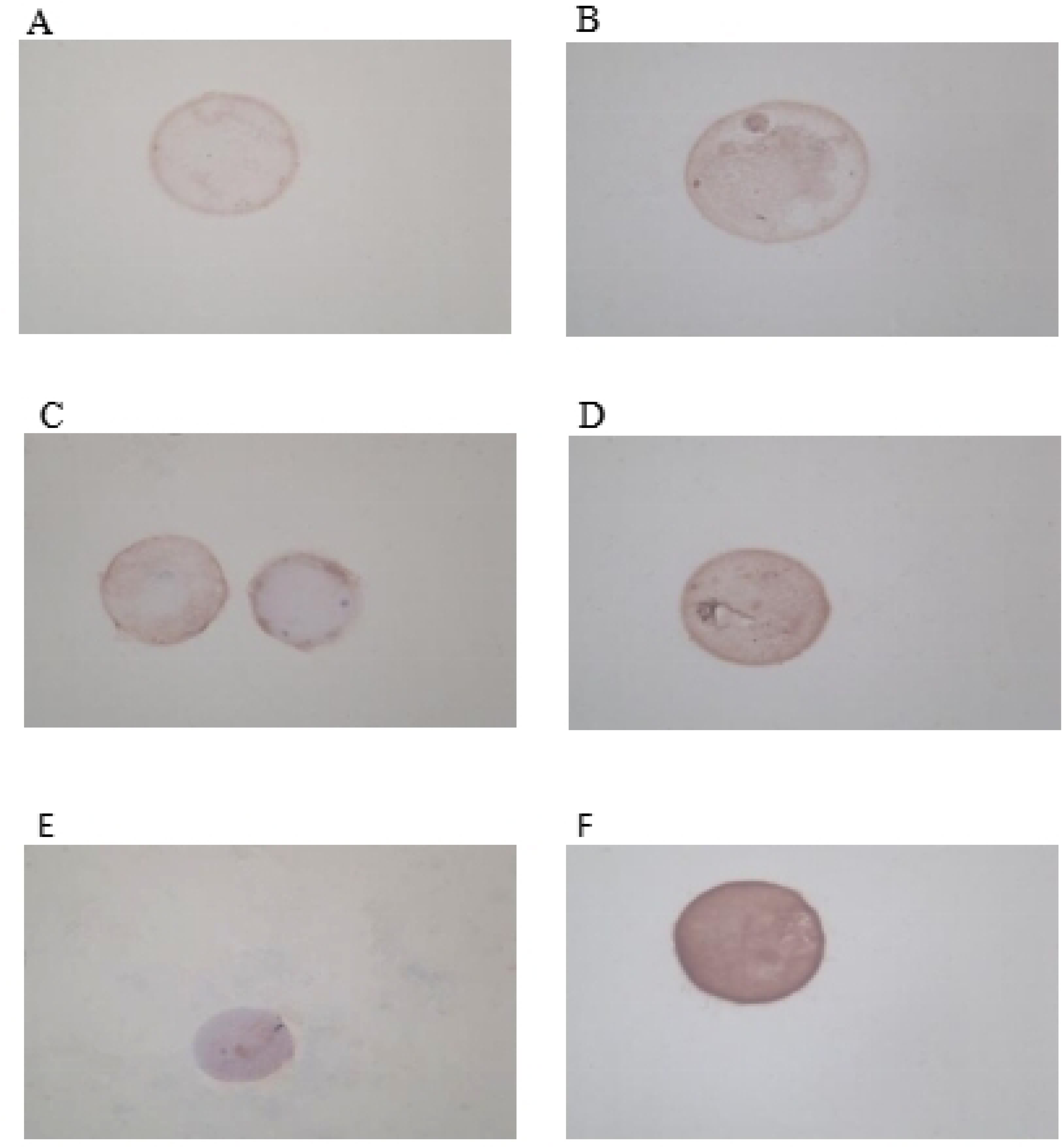
Expression of BMP-6 proten in occyte with immunoistochemistry(×400)A:Control group; B:COH group; C:Shugan low group; D:Shugan high group; E:Bushen low group; F:Bushen high group.

**Fig. 6.**
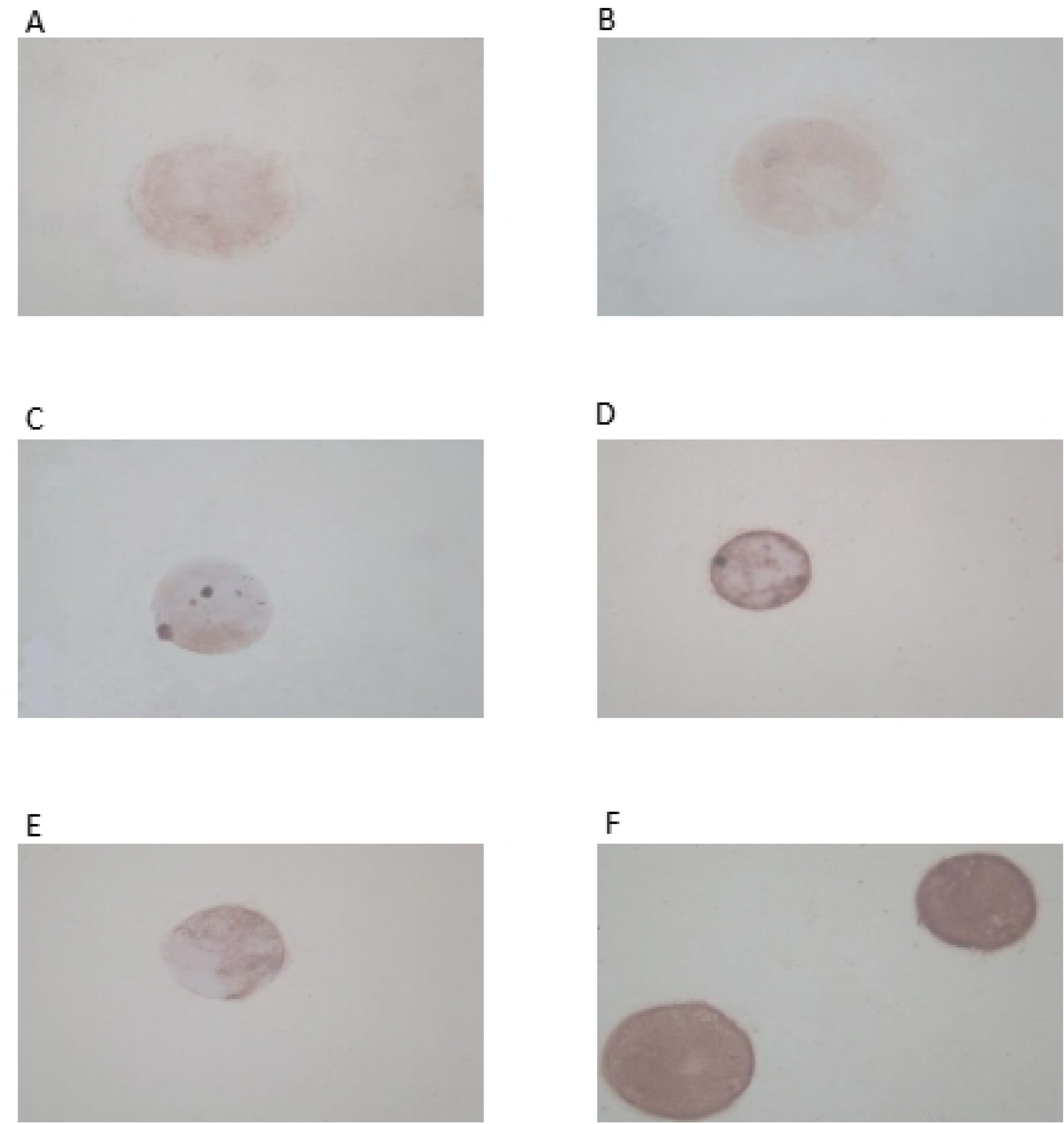
Expression of ALK-2 protein in occyte with immunohistochemistry(×400)A:Control group; B:COH group; C:Shugan low group; D:Shugan high group; E:Bushen low group; F:Bushen high group.

**Fig. 7.**
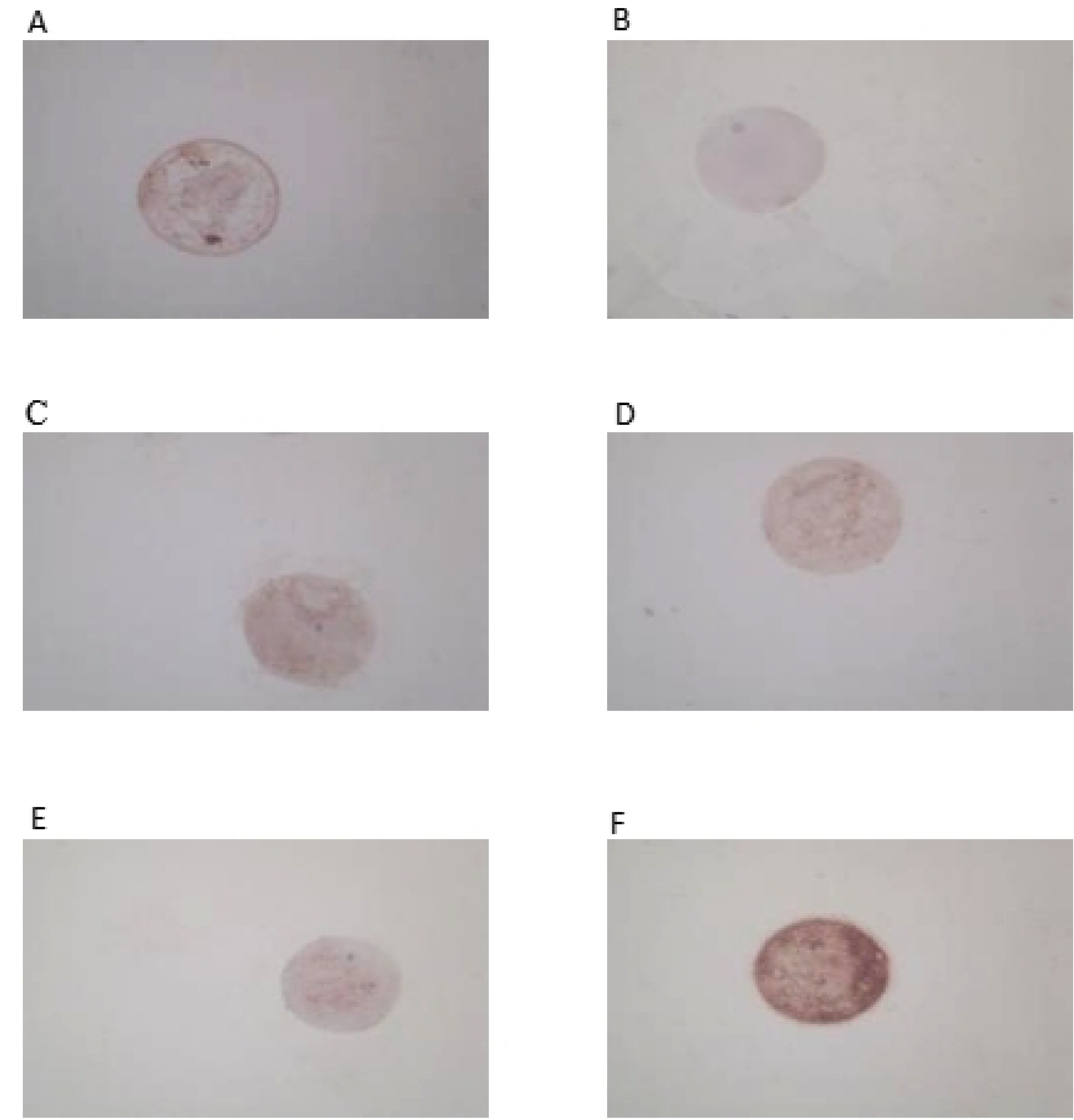
Expression of ALK-6 protein in occyte with immunohistochemistry(×400)A:Control group; B:COH group; C:Shugan low group; D:Shugan high group; E:Bushen low group; F:Bushen high group.

**Table 4.** BMP-6, ALK-2 and ALK-6 positive expression levels in the oocytes of each group (x ± S)

| Group | BMP-6 | ALK-2 | ALK-6 |
| --- | --- | --- | --- |
| Normal | 0.67±0.58□▽#▲ | 0.67±0.58□▽#▲ | 0.33±0.58□▽#▲ |
| Control | 1.00±1.00□▽#▲ | 1.33±0.58□▽#▲ | 0.67±0.58□▽#▲ |
| Shugan low | 3.67±0.58*△▽▲ | 3.00±1.00*△▽▲ | 2.67±0.58*△▽#▲ |
| Shugan high | 6.67±1.15*△□#▲ | 6.67±1.15*△□#▲ | 4.67±1.15*△□#▲ |
| Bushen low | 3.33±0.58*△▽▲ | 4.67±1.15*△▽▲ | 6.67±1.15*△□▽▲ |
| Bushen high | 8.67±0.58*△□▽# | 8.67±0.58*△□▽# | 8.67±0.58*△□▽# |
| Compared with Control group:*P<0.05; |  | Compared with COH group:△P<0.05; |  |
| Compared with shugan low group:□P<0.05; |  | Compared with shugan high group:▽P<0.05; |  |
| Compared with Bushen low group:P<0.05; |  | Compared with Bushen high group:▲P<0.05 |  |

#### 3.2.2 Comparison of Smad1/5 protein expression (Figure 8-9, Table 5)

Compared with the blank control group, the IHS score of the COH group was not statistically significant (P>0.05); Comparison between treatment groups: The IHS score of the high-dose kidney tonifying group was higher than the other three groups (P<0.05); The expression of kidney tonifying low-dose group was better than that of liver soothing high-dose group and liver soothing low-dose group (P<0.05); The IHS score of the high-dose group of liver soothing therapy increased compared to the low-dose group of liver soothing therapy (P<0.05).

**Fig. 8.**
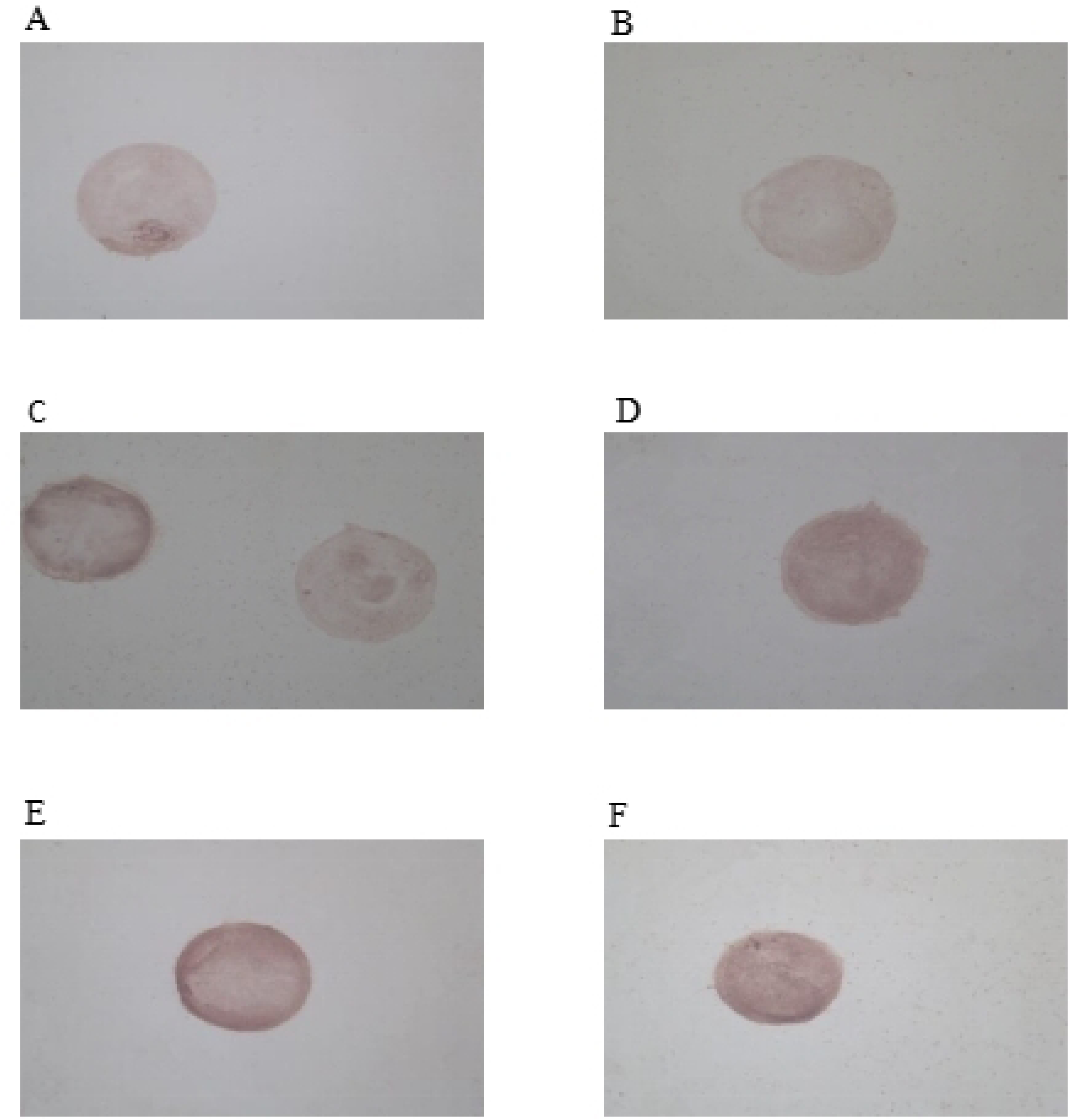
Expression of Smad1 protein in occyte with immunohistochemistry(×400)A:Control group; B:COH group; C:Shugan low group; D:Shugan high group; E:Bushen low group; F:Bushen high group.

**Fig. 9.**
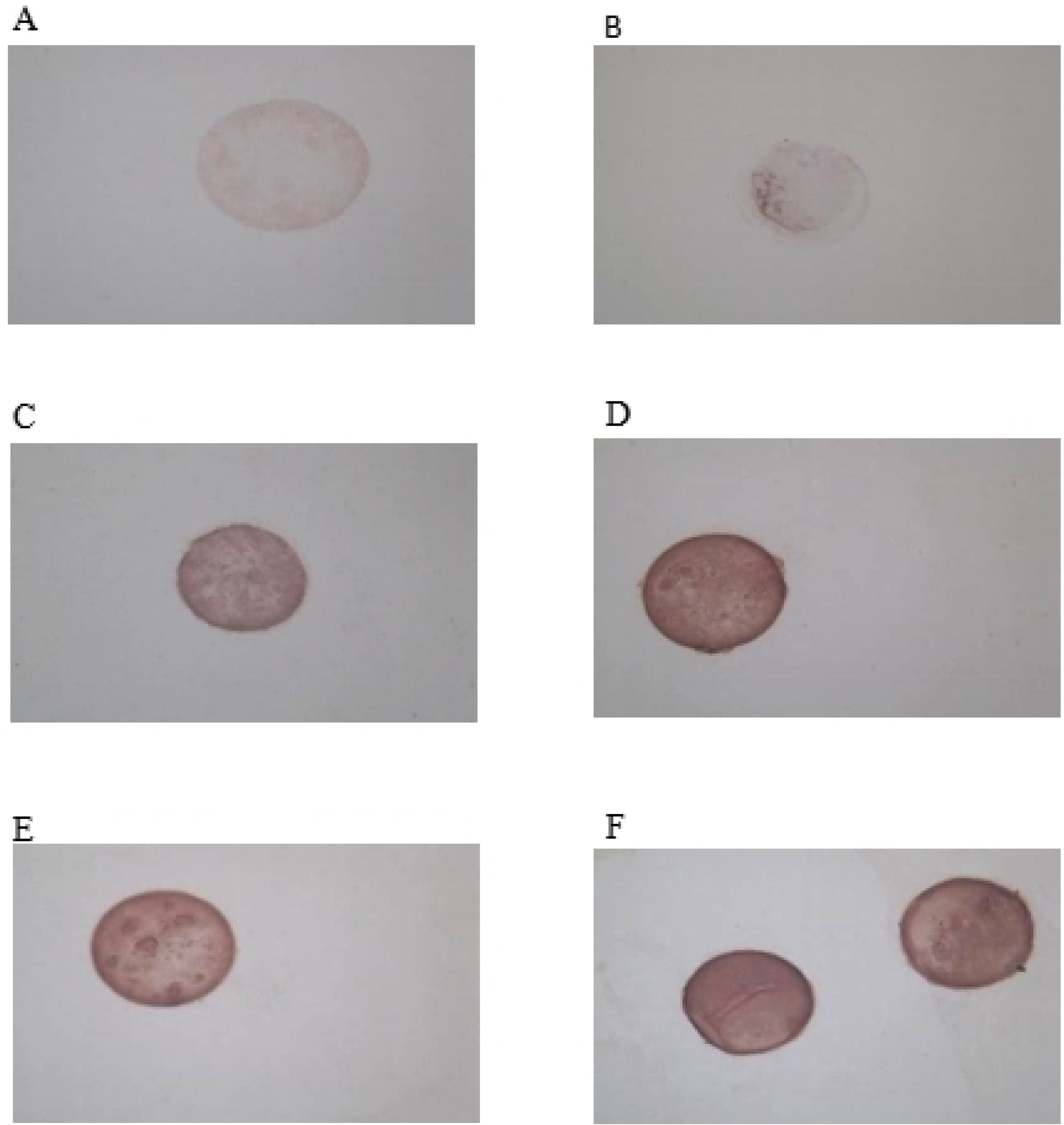
Expression of Smad5 protein in occyte with immunohistochemistry(×400)A:Control group; B:COH group; C:Shugan low group; D:Shugan high group; E:Bushen low group; F:Bushen high group.

**Table 5.** Smad1 and Smad5 positive expression levels in the oocytes of each group.

| Group | Smad1 | Smad5 |
| --- | --- | --- |
| Normal | 0.67±0.58□▽#▲ | 1.00±1.00□▽#▲ |
| Control | 0.67±1.15□▽#▲ | 0.33±0.58□▽#▲ |
| Shugan low | 2.67±0.58*Δ▽#▲ | 3.00±1.00*Δ▽#▲ |
| Shugan high | 4.67±1.15*Δ□#▲ | 8.67±0.58*Δ□#▲ |
| Bushen low | 6.67±1.15*Δ□▽▲ | 5.33±1.15*Δ□▽▲ |
| Bushen high | 8.67±0.58*Δ□▽# | 11.00±1.73*Δ□▽# |

Compared with Control group:\*P<0.05; Compared with COH group:ΔP<0.05;
Compared with shugan low group:□P<0.05; Compared with shugan high group:▽P<0.05;
Compared with Bushen low group:#P<0.05; Compared with Bushen high group:▲P<0.05

#### 3.2.3 Comparison of Smad8 protein expression (Figure 10, Table 6)

Compared with the blank control group, the expression rate of positive cells in the COH group decreased, and the IHS score was not statistically significant (P>0.05); The scores of each treatment group were higher than those of the blank control group and the COH group (P<0.05); The IHS score in Bushen high group was higher than that in the low dose group (P<0.05); The expression of high dose Shugan group was significantly higher than that of low dose Shugan group (P<0.05); There was no statistically significant difference between the high-dose groups (P>0.05); The IHS score in Bushen low group was higher than that in shugan low group (P<0.05).

**Fig. 10.**
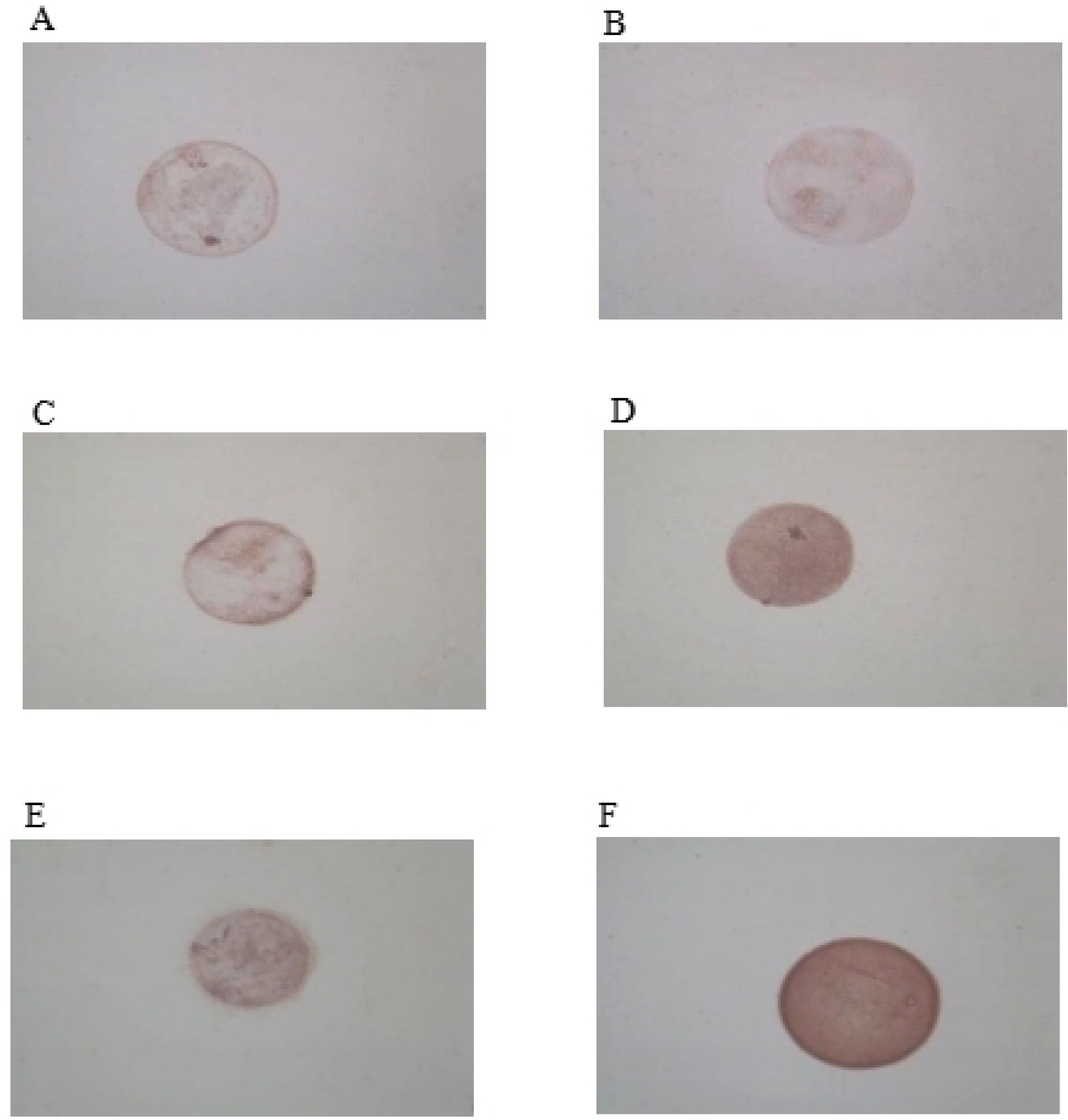
Expression of Smad8 protein in occyte with immunohistochemistry(×400)A:Normal group; B:Control group; C:Shugan low group; D:Shugan high group; E:Bushen low group; F:Bushen high group.

**Table 6.** Smad8 and Smad4 positive expression levels in the oocytes of each group.

| Group | Smad8 | Smad4 |
| --- | --- | --- |
| Normal | 0.67±0.58□▽#▲ | 1.00±1.00□▽#▲ |
| Control | 1.00±1.00□▽#▲ | 1.67±0.58□▽#▲ |
| Shugan low | 2.67±0.58*Δ▽#▲ | 4.33±1.53*Δ▽▲ |
| Shugan high | 8.67±0.58*Δ□# | 8.67±0.58*Δ□#▲ |
| Bushen low | 4.67±1.15*Δ□▽▲ | 4.67±1.15*Δ▽▲ |
| Bushen high | 8.33±0.58*Δ□# | 11.00±1.73*Δ□▽# |

Compared with Control group:\*P<0.05; Compared with COH group:ΔP<0.05;
Compared with shugan low group:□P<0.05; Compared with shugan high group:▽P<0.05;
Compared with Bushen low group:#P<0.05; Compared with Bushen high group:▲P<0.05

#### 3.2.4 Comparison of Smad4 protein expression (Figure 11, Table 6)

Compared with the blank control group, the IHS score in the COH group was not statistically significant (P>0.05); The scores of each treatment group were higher than those of the blank control group and the COH group (P<0.05); The IHS score in the high dose group was higher than that in the low dose group (P<0.05); The expression in Bushen high group was higher than that in shugan high group (P<0.05); There was no statistically significant difference between the low dose groups (P>0.05).

**Fig. 11.**
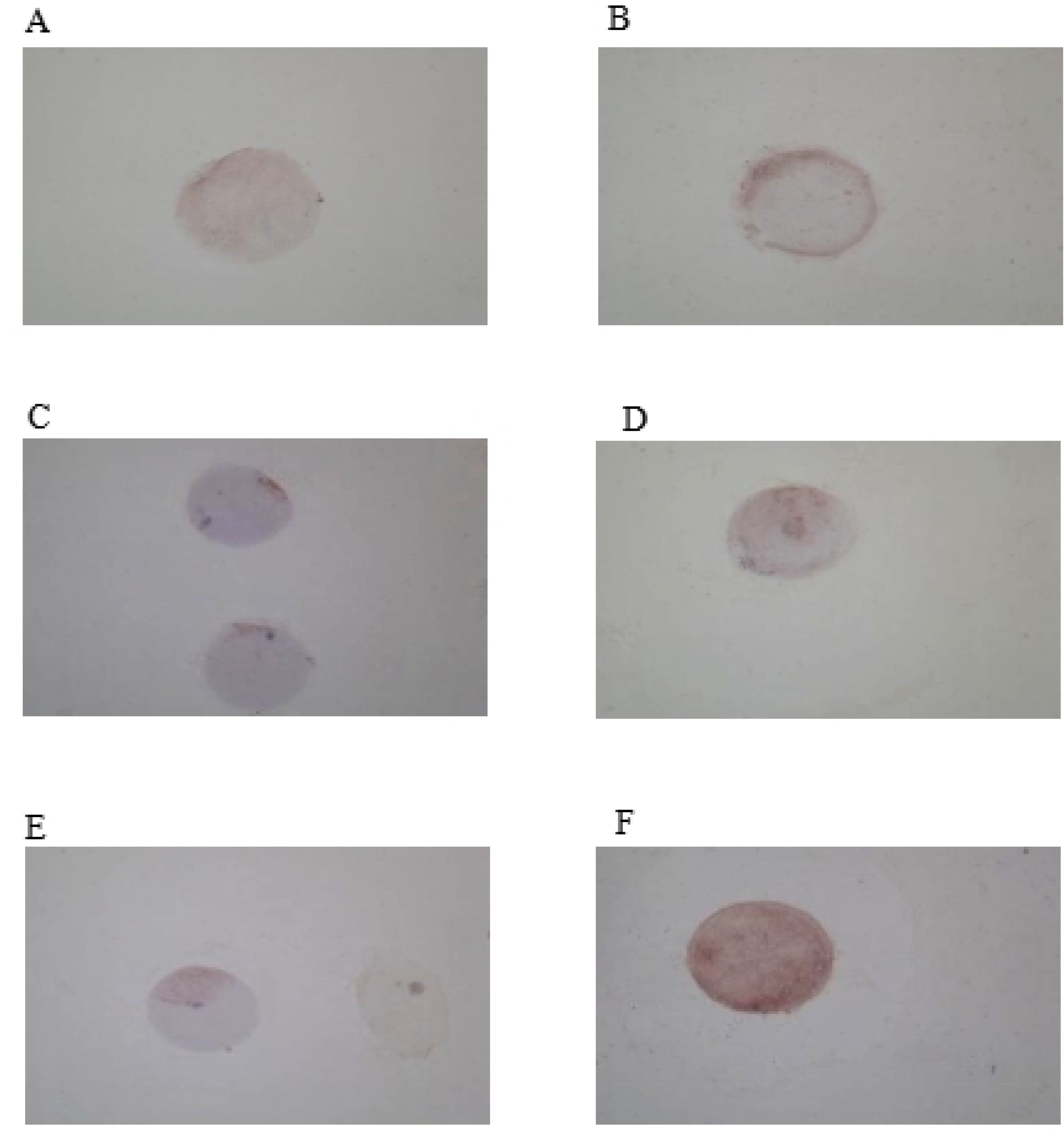
Expression of Smad4 protein in occyte with immunohistochemistry(×400)A:Control group; B:COH group; C:Shugan low group; D:Shugan high group; E:Bushen low group; F:Bushen high group.

### 3.3 Comparison of mRNA and protein expressions of BMP-6, ALK-2/6, Smad1/5/8/4 in mouse oocytes of each group detected by real-time fluorescence quantitative PCR and Western blot

#### 3.3.1 Comparison of BMP-6 mRNA expression and protein expression (Figure 12, Table 7)

Compared with the blank control group, the expression of BMP-6 mRNA in the COH group was not statistically significant (P>0.05), while the expression of BMP-6 protein decreased (P<0.05). The expression of BMP-6 mRNA in each treatment group increased compared to the blank control group and COH group (P<0.05); Comparison between treatment groups: The expression of BMP-6 in the high-dose group of kidney tonifying therapy was higher than that in the other three groups (P<0.05); The expression of BMP-6 in the high-dose group of liver soothing was higher than that in the low-dose group (P<0.05); There was no statistically significant difference (P>0.05) in the comparison between low-dose groups.

**Fig. 12.**
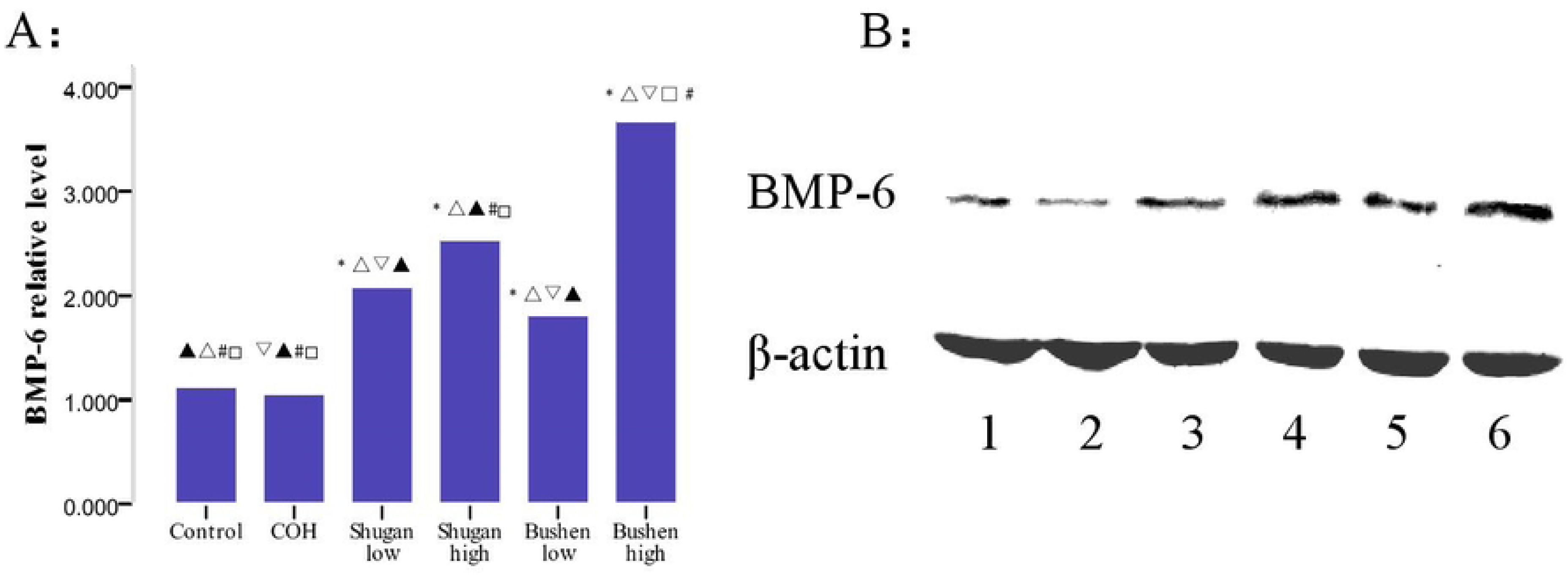
Expression of BMP-6 in the mice oocytes A: *P<0.05,compared with Control group;△P<0.05,compared with COH group; □P,0.05,compared with shugan low group;▽P<0.05,compared with shugan high group;#P<0.05,compared with Bushen low group; ▴P<0.05,compared with Bushen high group B: 1 Control group; 2 COH group; 3 Shugan low group; 4 Shugan high group; 5 Bushen low group; 6 Bushen high group.

**Table 7.** Expression of BMP-6 m RNA and protein of the oocyte in the mice (x ± S)

| Group | BMP-6 mRNA | BMP-6 protein |
| --- | --- | --- |
| Normal | 1.125±0.333□▽#▲ | 0.161±0.003□▽#▲ |
| Control | 1.058±0.176□▽#▲ | 0.160±0.004□▽#▲ |
| Shugan low | 2.090±0.172*Δ▽▲ | 0.213±0.004*Δ▽▲ |
| Shugan high | 2.544±0.245*Δ□#▲ | 0.302±0.007*Δ□#▲ |
| Bushen low | 1.821±0.108*Δ▽▲ | 0.219±0.009*Δ▽▲ |
| Bushen high | 3.679±0.232*Δ□▽# | 0.340±0.006*Δ□▽# |

Compared with Control group:\*P<0.05;
Compared with COH group:ΔP<0.05;
Compared with shugan low group:□P<0.05;
Compared with shugan high group:▽P<0.05;
Compared with Bushen low group:#P<0.05;
Compared with Bushen high group:▲P<0.05

#### 3.3.2 Comparison of ALK-2 mRNA expression and protein expression (Figure 13, Table 8)

Compared with the blank control group, the expression of ALK-2 in the COH group was not statistically significant (P>0.05); Inter group comparison showed that the expression of ALK-2 mRNA in the high-dose group of liver soothing was higher than that in the low-dose group of kidney tonifying and low-dose group of liver soothing (P<0.05); The expression of ALK-2 protein in the high-dose group was higher than that in the low-dose group (P<0.05), and the expression in the high-dose group of kidney tonifying was enhanced compared to the high-dose group of liver soothing (P<0.05); The expression of ALK-2 mRNA and protein between low-dose groups was not statistically significant (P>0.05).

**Fig. 13.**
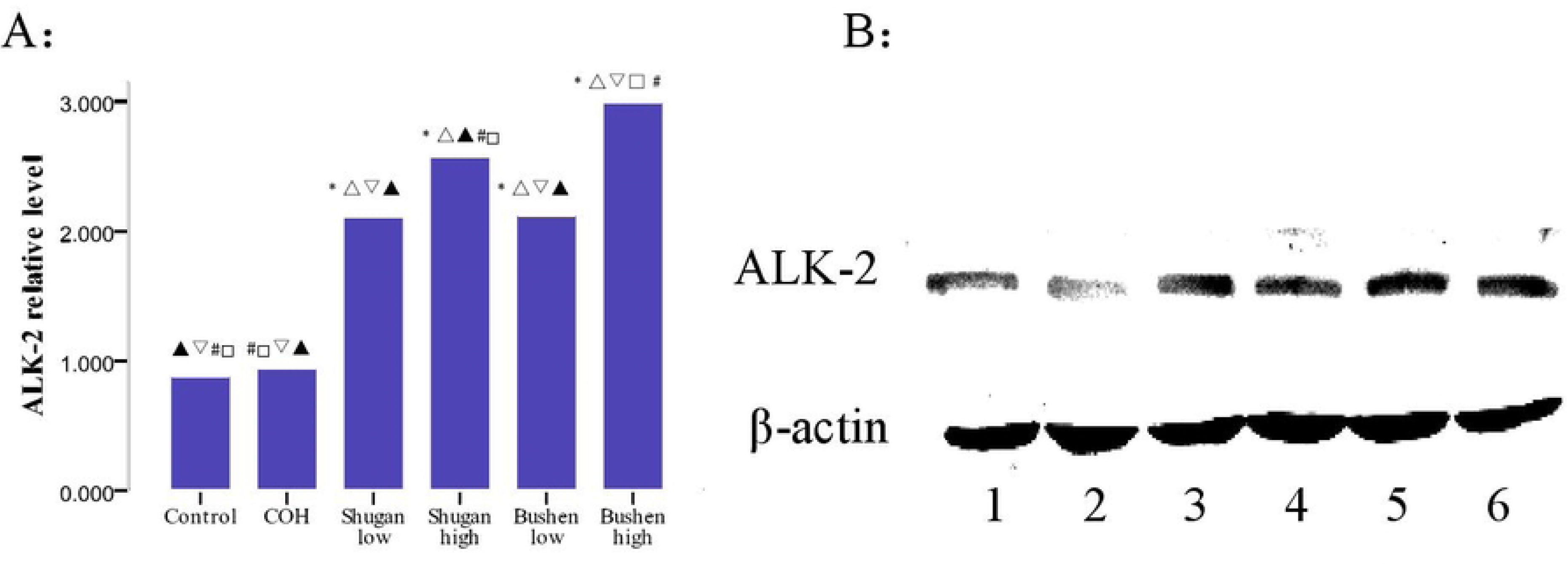
Expression of ALK-2 in the mice oocytes A: *P<0.05,compared with Control group;△P<0.05,compared with COH group; □P,0.05,compared with shugan low group;▽P<0.05,compared with shugan high group;#P<0.05,compared with Bushen low group; ▴P<0.05,compared with Bushen high group B: 1 Control group; 2 COH group; 3 Shugan low group; 4 Shugan high group; 5 Bushen low group; 6 Bushen high group.

**Table 8.**
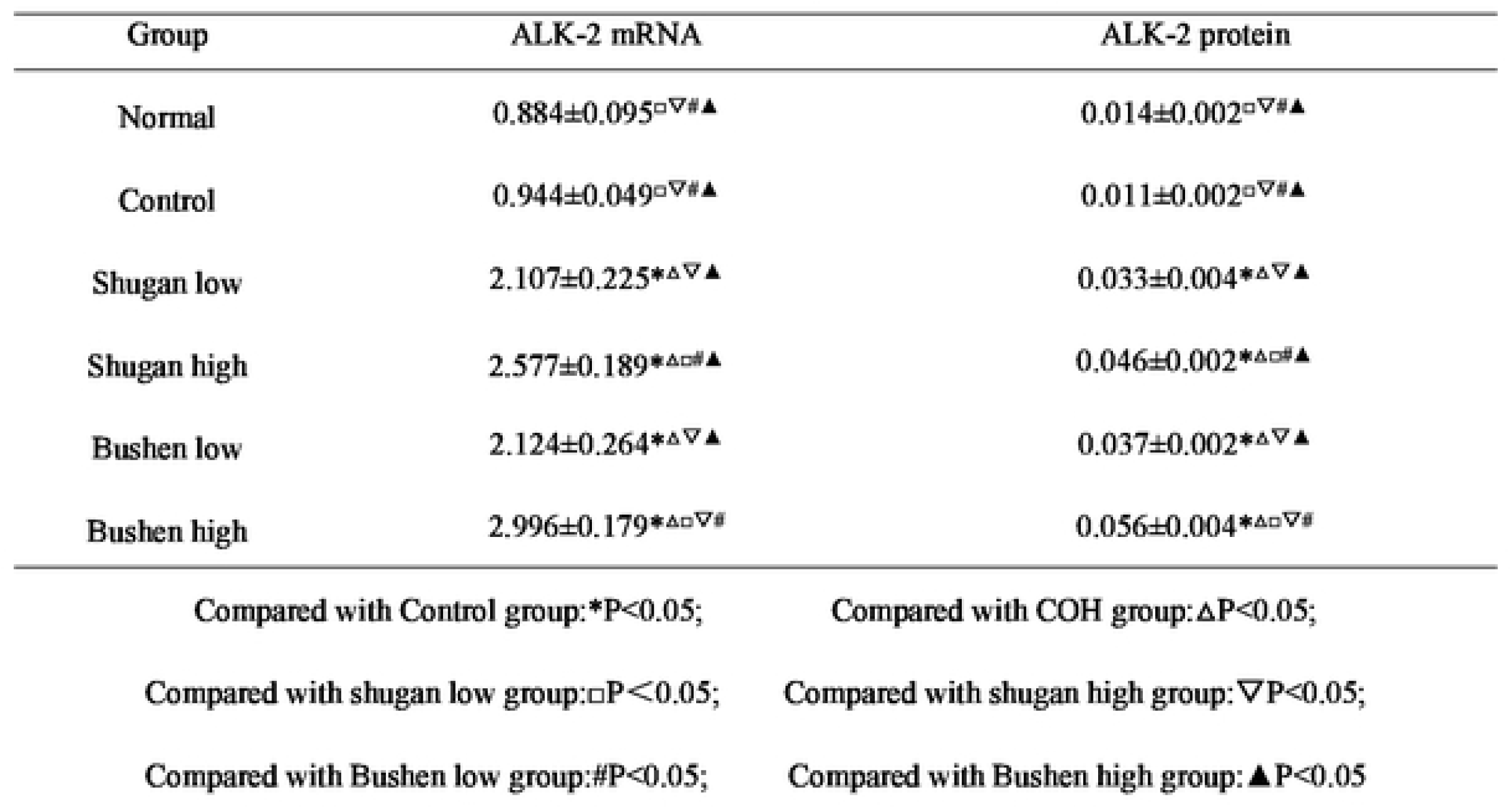
Expression of ALK-2 m RNA and protein of the oocyte in the mice (x ± S)

| Group | ALK-2 mRNA | ALK-2 protein |
| --- | --- | --- |
| Normal | 0.884±0.095□▽#▲ | 0.014±0.002□▽#▲ |
| Control | 0.944±0.049□▽#▲ | 0.011±0.002□▽#▲ |
| Shugan low | 2.107±0.225*Δ▽▲ | 0.033±0.004*Δ▽▲ |
| Shugan high | 2.577±0.189*Δ□#▲ | 0.046±0.002*Δ□#▲ |
| Bushen low | 2.124±0.264*Δ▽▲ | 0.037±0.002*Δ▽▲ |
| Bushen high | 2.996±0.179*Δ□▽# | 0.056±0.004*Δ□▽# |

Compared with Control group:\*P<0.05;
Compared with COH group:ΔP<0.05;
Compared with shugan low group:□P<0.05;
Compared with shugan high group:▽P<0.05;
Compared with Bushen low group:#P<0.05;
Compared with Bushen high group:▲P<0.05

#### 3.3.3 Comparison of ALK-6 mRNA expression and protein expression (Figure 14, Table 9)

Compared with the blank control group, the expression of ALK-6 in the COH group was not statistically significant (P>0.05); The expression of ALK-6 mRNA and protein was increased in each treatment group compared to the blank control group and COH group (P<0.05); The expression of kidney tonifying high-dose group was higher than that of the other three groups (P<0.05); The expression of kidney tonifying low-dose group was higher than that of liver soothing group (P<0.05); The expression of ALK-6 mRNA in the high-dose group of Shugan was enhanced compared to the low-dose group of Shugan (P<0.05).

**Fig. 14.**
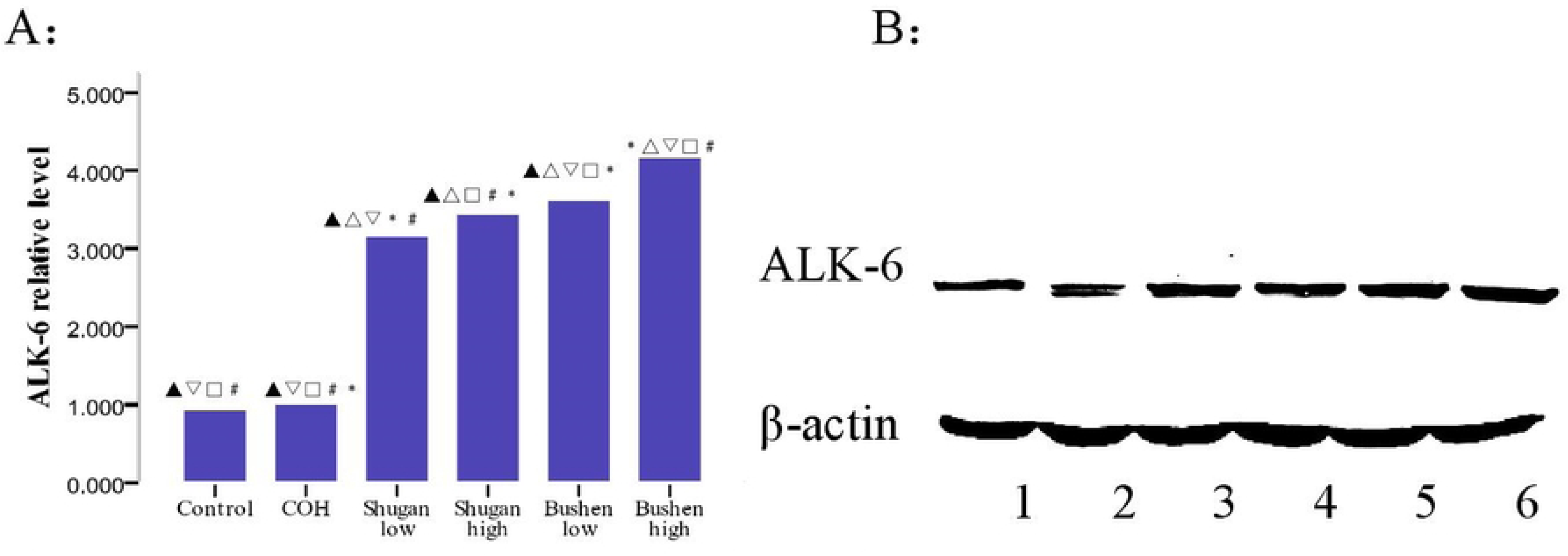
Expression of ALK-6 in the mice oocytes A: *P<0.05,compared with Control group;△P<0.05,compared with COH group; □P,0.05,compared with shugan low group;▽P<0.05,compared with shugan high group;#P<0.05,compared with Bushen low group; ▴P<0.05,compared with Bushen high group B: 1 Control group; 2 COH group; 3 Shugan low group; 4 Shugan high group; 5 Bushen low group;

**Table 9.**
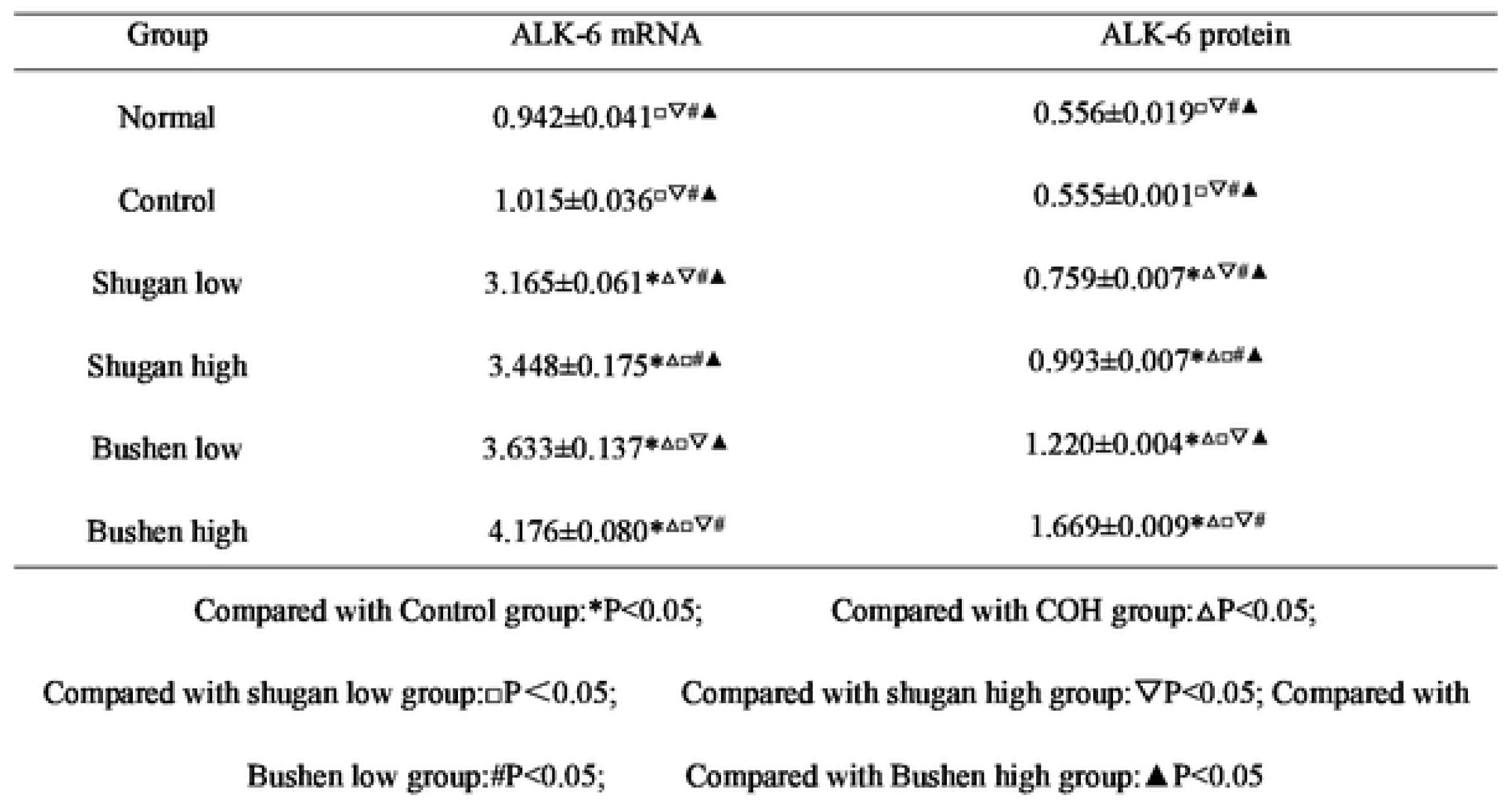
Expression of ALK-6 m RNA and protein of the oocyte in the mice (x ± S)

**Table 10.** Expression of Smad1 m RNA and protein of the oocyte in the mice (x ± S)

| Table10 Expression of Smad1 m RNA and protein of the oocyte in the mice ( $\bar{x} \pm S$ ) | | |
| --- | --- | --- |
| Group | Smad1 mRNA | Smad1 protein |
| Normal | 1.063±0.055$\square\nabla\#\blacktriangle$ | 0.290±0.011$\square\nabla\#\blacktriangle$ |
| Control | 1.084±0.073$\square\nabla\#\blacktriangle$ | 0.270±0.014$\square\nabla\#\blacktriangle$ |
| Shugan low | 2.393±0.160$\ast\Delta\nabla\#\blacktriangle$ | 0.314±0.009$\ast\Delta\nabla\#\blacktriangle$ |
| Shugan high | 2.734±0.270$\ast\Delta\square\#\blacktriangle$ | 0.466±0.020$\ast\Delta\square\#\blacktriangle$ |
| Bushen low | 3.358±0.225$\ast\Delta\square\nabla\blacktriangle$ | 0.560±0.003$\ast\Delta\square\nabla\blacktriangle$ |
| Bushen high | 3.870±0.168$\ast\Delta\square\nabla\#$ | 0.715±0.012$\ast\Delta\square\nabla\#$ |
| Compared with Control group:$\ast P<0.05$;<br>Compared with shugan low group:$\square P<0.05$;<br>Compared with Bushen low group:$\# P<0.05$;<br>Compared with COH group:$\Delta P<0.05$;<br>Compared with shugan high group:$\nabla P<0.05$;<br>Compared with Bushen high group:$\blacktriangle P<0.05$ |  |  |

#### 3.3.4 Comparison of Smad1 mRNA expression and protein expression (Figure 15, Table10)

Compared with the blank control group, the expression of Smad1 in the COH group was not statistically significant (P>0.05); The expression levels increased in the kidney tonifying group and liver soothing group (P<0.05); Compared with the high-dose group of kidney tonifying, the low-dose group of kidney tonifying and the high-dose group of liver soothing showed lower expression (P<0.05); The expression of kidney tonifying low-dose group was better than that of liver soothing group (P<0.05); The expression of Smad1 increased in the high-dose group of liver soothing compared to the low-dose group (P<0.05).

**Fig. 15.**
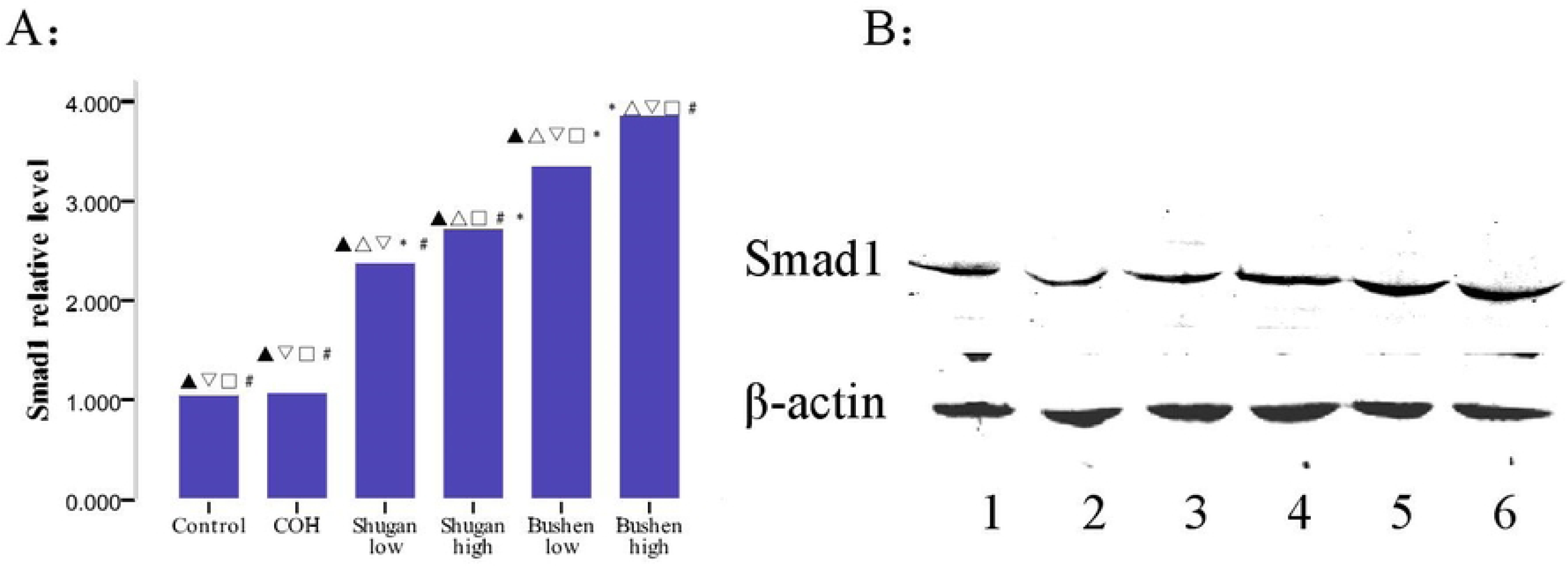
Expression of Smad1 in the mice oocytes A: *P<0.05,compared with Control group;△P<0.05,compared with COH group; □P,0.05,compared with shugan low group;▽P<0.05,compared with shugan high group;#P<0.05,compared with Bushen low group; ▴P<0.05,compared with Bushen high group B: 1 Control group; 2 COH group; 3 Shugan low group; 4 Shugan high group; 5 Bushen low group; 6 Bushen high group.

#### 3.3.5 Comparison of Smad5 mRNA expression and protein expression (Figure 16, Table 11)

Compared with the blank control group, the expression of Smad5 in the COH group was not statistically significant (P>0.05); Compared with the high-dose group of kidney tonifying, the low-dose group of kidney tonifying and the high-dose group of liver soothing showed a decrease in expression (P<0.05); The expression of Smad5 in the high-dose group of liver soothing therapy was higher than that in the low-dose group of kidney tonifying therapy and the low-dose group of liver soothing therapy (P<0.05); Compared with the low-dose group of liver soothing and kidney tonifying, the expression of Smad5 was reduced (P<0.05).

**Fig. 16.**
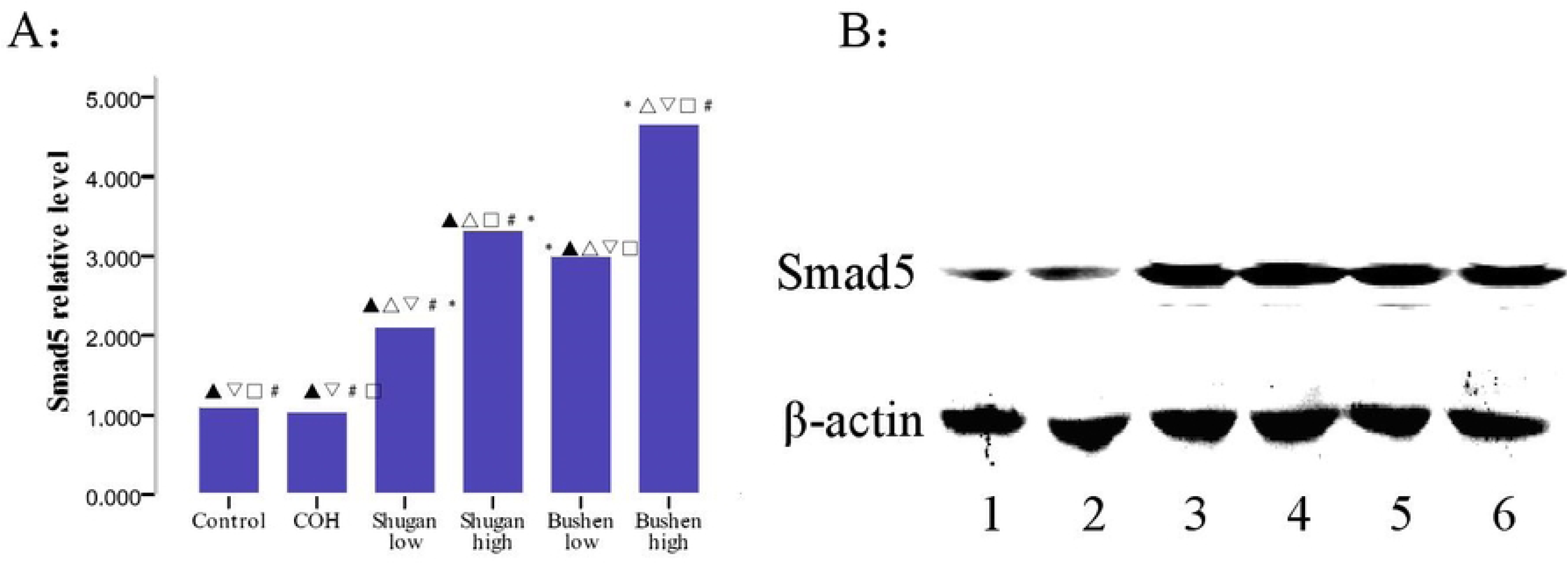
Expression of Smad5 in the mice oocytes A: *P<0.05,compared with Control group;△P<0.05,compared with COH group; □P,0.05,compared with shugan low group;▽P<0.05,compared with shugan high group;#P<0.05,compared with Bushen low group; ▴P<0.05,compared with Bushen high group B: 1 Control group; 2 COH group; 3 Shugan low group; 4 Shugan high group; 5 Bushen low group; 6 Bushen high group.

**Table 11.**
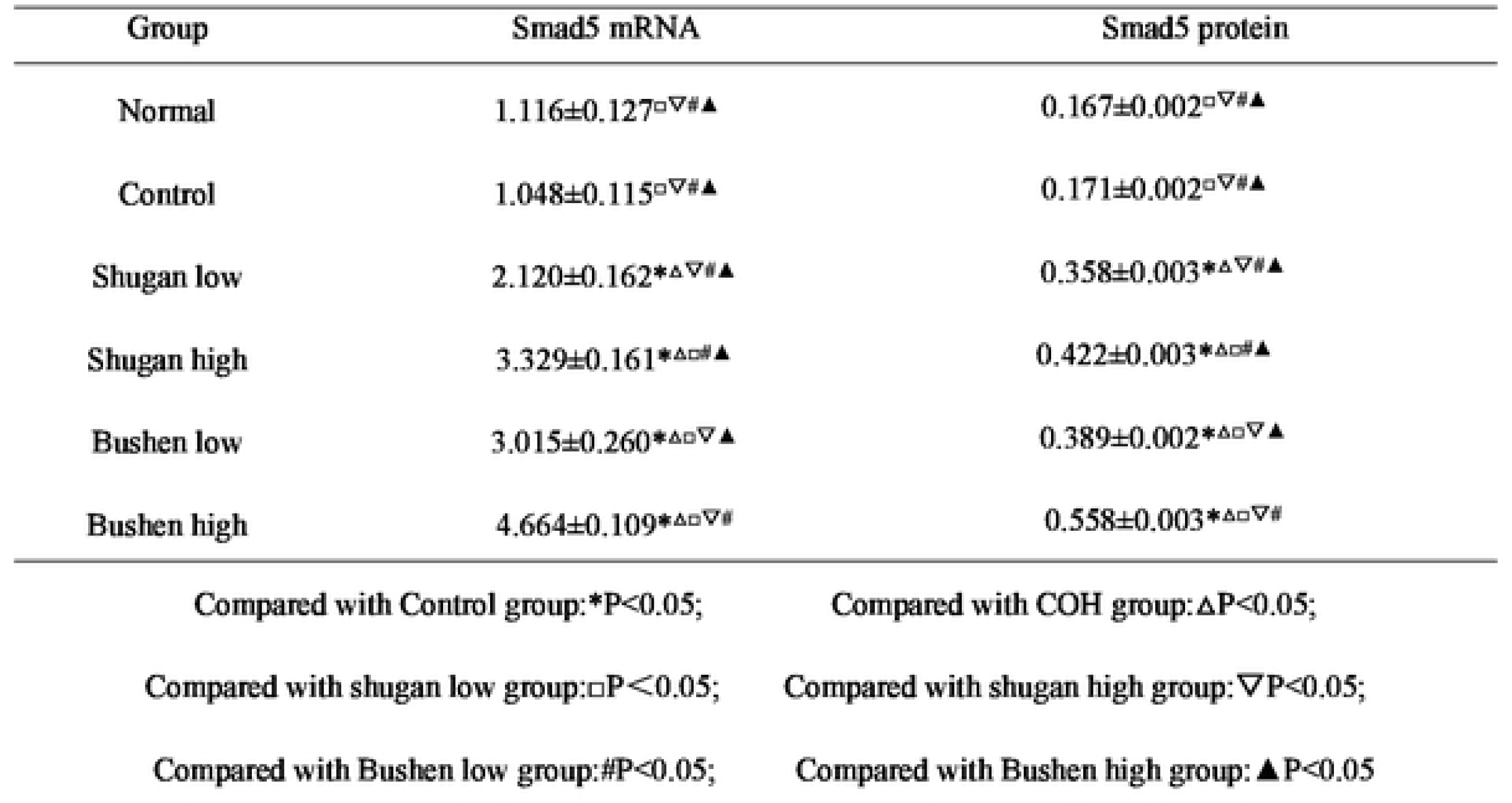
Expression of Smad5 m RNA and protein of the oocyte in the mice (x ± S)

| Group | Smad5 mRNA | Smad5 protein |
| --- | --- | --- |
| Normal | 1.116±0.127□▽#▲ | 0.167±0.002□▽#▲ |
| Control | 1.048±0.115□▽#▲ | 0.171±0.002□▽#▲ |
| Shugan low | 2.120±0.162*Δ▽#▲ | 0.358±0.003*Δ▽#▲ |
| Shugan high | 3.329±0.161*Δ□#▲ | 0.422±0.003*Δ□#▲ |
| Bushen low | 3.015±0.260*Δ□▽▲ | 0.389±0.002*Δ□▽▲ |
| Bushen high | 4.664±0.109*Δ□▽# | 0.558±0.003*Δ□▽# |

Compared with Control group:\*P<0.05;
Compared with COH group:ΔP<0.05;
Compared with shugan low group:□P<0.05;
Compared with shugan high group:▽P<0.05;
Compared with Bushen low group:#P<0.05;
Compared with Bushen high group:▲P<0.05

#### 3.3.6 Comparison of Smad8 mRNA expression and protein expression (Figure 17, Table 12)

Compared with the blank control group, the expression of COH group was not statistically significant (P>0.05); The expression of Smad8 in each treatment group was significantly higher than that in the blank control group and COH group (P<0.05); Inter group comparison showed that the expression in the high-dose group increased compared to the low-dose group (P<0.05); There was no significant difference (P>0.05) between the high-dose group for tonifying the kidney and the high-dose group for soothing the liver; The expression of Smad8 mRNA in the low-dose group of tonifying the kidney was better than that in the low-dose group of soothing the liver (P<0.05).

**Fig. 17.**
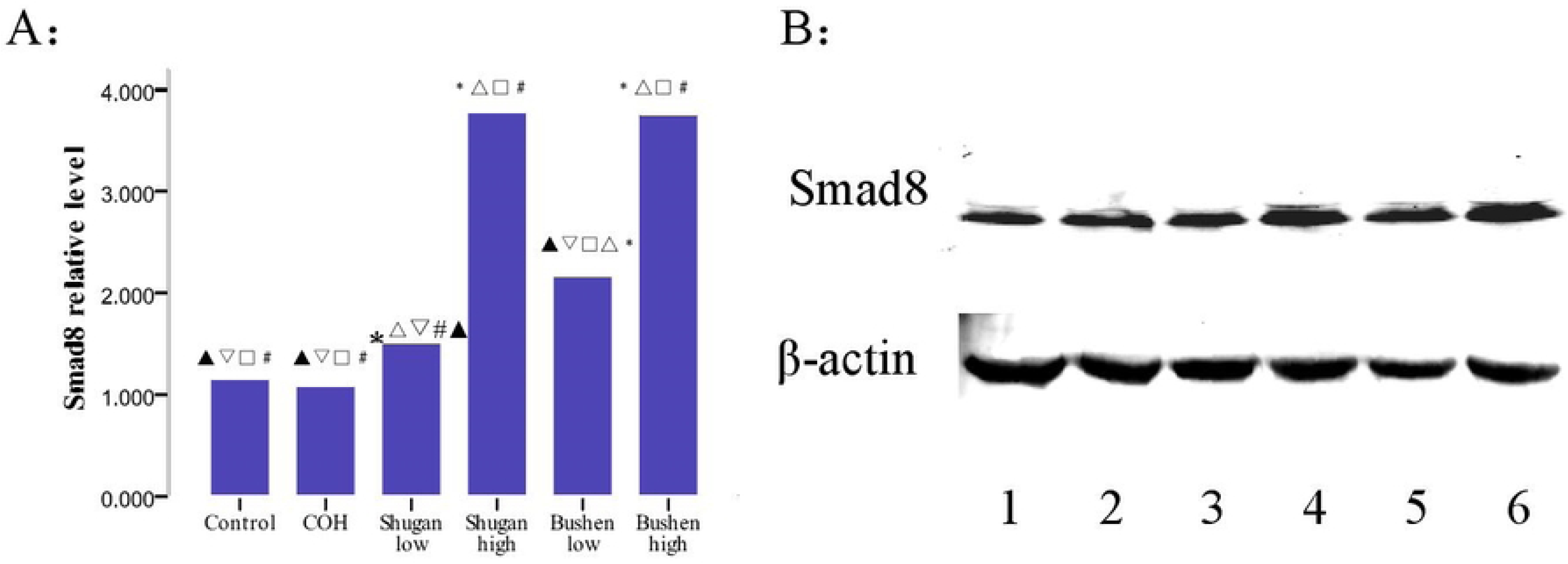
Expression of Smad8 in the mice oocytes A: *P<0.05,compared with Control group;△P<0.05,compared with COH group; □P,0.05,compared with shugan low group;▽P<0.05,compared with shugan high group;#P<0.05,compared with Bushen low group; ▴P<0.05,compared with Bushen high group B: 1 Control group; 2 COH group; 3 Shugan low group; 4 Shugan high group; 5 Bushen low group; 6 Bushen high group.

**Table 12.** Expression of Smad8 m RNA and protein of the oocyte in the mice (x ± S)

|  |  |  |
| --- | --- | --- |
| Group | Smad8 mRNA | Smad8 protein |
| Normal | 1.159±0.139□▽#▲ | 0.428±0.004□▽#▲ |
| Control | 1.088±0.134□▽#▲ | 0.421±0.008□▽#▲ |
| Shugan high | 3.780±0.167*△□# | 1.033±0.004*△□# |
| Bushen low | 2.173±0.179*△□▽▲ | 0.727±0.006*△□▽▲ |
| Bushen high | 3.763±0.409*△□# | 1.035±0.002*□# |
| Compared with Control group:*P<0.05;<br>Compared with shugan low group:□P<0.05;<br>Compared with Bushen low group:#P<0.05;<br>Compared with COH group:△P<0.05;<br>Compared with shugan high group:▽P<0.05;<br>Compared with Bushen high group:▲P<0.05 |  |  |

#### 3.3.7 Comparison of Smad4 mRNA expression and protein expression (Figure 18, Table 13)

Compared with the blank control group, the expression of COH group was not statistically significant (P>0.05); The expression of Smad4 in each treatment group was significantly higher than that in the blank control group and COH group (P<0.05); Inter group comparison showed that the expression in the high-dose group was enhanced compared to the low-dose group (P<0.05); There was no statistically significant difference (P>0.05) between the low-dose group for tonifying the kidney and the low-dose group for soothing the liver; The expression of Smad4 in the high-dose group of tonifying the kidney was better than that in the low-dose group of soothing the liver (P<0.05)

**Fig. 18.**
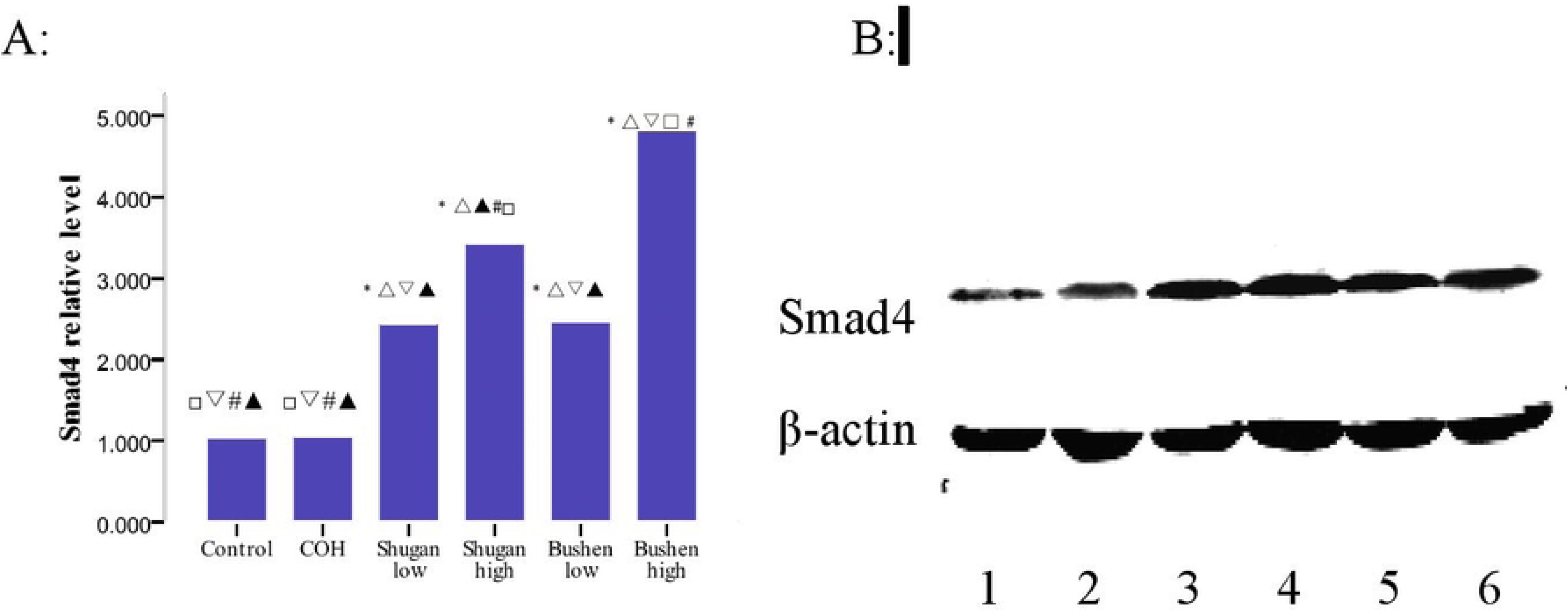
Expression of the Smad4 in the mice oocytes A: *P<0.05,compared with Control group;△P<0.05,compared with COH group; □P,0.05,compared with shugan low group;▽P<0.05,compared with shugan high group;#P<0.05,compared with Bushen low group; ▴P<0.05,compared with Bushen high group B: 1 Control group; 2 COH group; 3 Shugan low group; 4 Shugan high group; 5 Bushen low group; 6 Bushen high group.

**Table 13.** Expression of Smad4 m RNA and protein of the oocyte in the mice (x ± S)

| Group | Smad4 mRNA | Smad4 protein |
| --- | --- | --- |
| Normal | 1.045±0.053□▽#▲ | 0.311±0.004□▽#▲ |
| Control | 1.063±0.094□▽#▲ | 0.307±0.002□▽#▲ |
| Shugan low | 2.450±0.113*Δ▽▲ | 0.595±0.006*Δ▽▲ |
| Shugan high | 3.428±0.046*Δ□#▲ | 0.633±0.005*Δ□#▲ |
| Bushen low | 2.464±0.126*Δ▽▲ | 0.603±0.002*Δ▽▲ |
| Bushen high | 4.823±0.081*Δ□▽# | 0.692±0.008*Δ□▽# |

Compared with Control group:\*P<0.05; Compared with COH group:ΔP<0.05;
Compared with shugan low group:□P<0.05; Compared with shugan high group:▽P<0.05;
Compared with Bushen low group:#P<0.05; Compared with Bushen high group:▲P<0.05

## 4. Discussion

The ovaries have reproductive and endocrine functions, and their dysfunction can lead to various gynecological endocrine diseases such as menstrual disorders, dysfunctional uterine bleeding, amenorrhea, polycystic ovary syndrome, and infertility. Therefore, the normal development and elimination of oocytes are crucial for human reproduction. Research has found that bone morphogenetic protein-6[11] (BMP-6), a paracrine signal secreted by oocytes, plays an autocrine or paracrine regulatory role in the ovaries, regulating follicular development [12–14], inducing bone development and formation, and regulating mammalian growth, cell differentiation, and apoptosis. Receptors that can interact with BMP-6 are present in most species (including humans) of oocytes that grow follicles Granular cells and membrane cells [15–19]. The BMP-6 signal is transduced through various type I and type II receptors. At present, ALK2, ALK3, and ALK6 have been identified as potential type I receptors for BMP-6. Among them, ALK2 and ALK6 have the strongest affinity for BMP-6, and their receptor complexes mainly transmit signals through Smad5 [20–22]. BMPR II, ActR – Ⅱ A, and ActR – Ⅱ B have been identified as potential type II receptors. However, the differences in BMP-6 receptors among different types of ovarian cells have not yet been determined [23]. Smads are transforming growth factors-β TGF-β) The downstream signal transduction molecules of the superfamily can transmit extracellular signals from the cell membrane to the nucleus, and in TGF-β Plays an important role in signal transduction. BMPs can bind to cell surface receptors to form ligand activated receptor complexes, which then activate downstream Smads signaling molecules through phosphorylation, thereby achieving the transmission of BMP signals from extracellular to intracellular target genes.

Traditional Chinese medicine believes that the kidneys store essence, providing a material basis for the development and maturation of follicles, ovulation, and the onset of menstruation; The liver stores blood and is responsible for regulating blood flow, which can regulate the sealing and opening of the kidneys, promote smooth flow of qi and blood, facilitate timely menstruation, and facilitate the smooth discharge of oocytes. The liver and kidney have two organs, one for storing and the other for releasing. They regulate the qi and blood of Tiangui and Chongren, and are closely related to the timely onset of menstruation and the normal development and discharge of oocytes.

The Kidney Tonifying and Meridian Regulating Formula (a basic formula for clinical treatment of menstrual disorders, infertility and other diseases caused by kidney deficiency, with the functions of tonifying the kidney and filling essence, regulating menstruation and aiding pregnancy) and Xiaoyao Pill (Xiaoyao Pill is a commonly used formula for harmonizing the liver and spleen, with the functions of soothing the liver and relieving depression, strengthening the spleen and regulating menstruation). The results of this experiment show that both can increase the expression of BMP-6 in the oocytes of mice in the treatment group, And successfully recruited their type I receptors ALK-2 and ALK-6 to activate them, phosphorylating their downstream signal receptors Smad1, Smad5, and Smad8, and then binding to Smad4 to initiate signal transduction. Confirming that the Kidney Tonifying and Meridian Regulating Formula and Xiaoyao Pill may affect the development of follicles and the quality of oocytes through BMP-6 and its receptors and downstream proteins, the Kidney Tonifying Method is more effective than the Liver Soothing Method. There may be differences in the mechanism of activating the Smads signaling pathway between the two methods. The tonifying kidney and regulating meridians formula may recruit ALK-2/6 by activating other type II receptors of BMP-6, further activating downstream Smads proteins, and thus initiating the transcription of target genes. In this process, the effect of the tonifying kidney high-dose group is significantly better than that of the low-dose group; And Xiaoyao Pill mainly recruits BMP-6 type II receptor BMPR II to form receptor complexes, and then recruits type I receptors ALK-2 and ALK-6, which may cause phosphorylation and activation. The activated type I receptor further phosphorylates intracellular signals Smad1/5/8, and then binds to Smad4 to initiate signal transduction. In this process, the effect of the high-dose group of liver soothing is significantly stronger than that of the low-dose group of liver soothing. In the transmission process of the entire BMP-6/Smads signaling pathway, the high-dose group of Bushen Tiaojing Formula was superior to the Shugan and low-dose groups in regulating BMP-6, ALK-2/6, and Smad1/5/4, while the low-dose group of Bushen Tiaojing Formula was superior to the Shugan group in regulating ALK6 and Smad1. The high-dose group of Shugan focused on regulating the expression of BMP-6, ALK-2, and Smad5, while there was no significant difference in activating Smad8 between the high-dose group of Bushen Tiaojing Formula and the high-dose group of Shugan. Therefore, the mechanisms of action of tonifying the kidney and soothing the liver each have their own emphasis, both of which can induce the transmission of Smads signaling pathway, further confirming the theory of “liver kidney homology”, “essence blood exchange”, “liver governing blood storage”, and “kidney governing reproduction” in traditional Chinese medicine. From the perspective of signaling pathway, it is confirmed that tonifying the kidney and soothing the liver have significant effects in intervening in follicular development and oocyte quality, This provides a solid experimental foundation for further exploration and clinical use as an adjuvant therapy for IVF-ET in the future.

## The author’s contribution

ZRJ and YLY make a significant contribution to conception and design, data acquisition, data analysis and interpretation and participation in drafting substantive revision of the manuscript or important content of the manuscript; JR make a significant contribution to conception and design, data acquisition, data analysis and interpretation; BJ participation in drafting substantive revision of the manuscript or important content of the manuscrip; DHL agree o be responsible for all aspects of the work and ensure that the accuracy or completeness of the work is properly investigated and resolved.

Approve the publication of the final version. Every author should fully participate And work.

